# Metacognition and the effect of incentive motivation in two compulsive disorders: gambling disorder and obsessive-compulsive disorder

**DOI:** 10.1101/2021.09.30.462582

**Authors:** Monja Hoven, Nina S. de Boer, Anna E. Goudriaan, Damiaan Denys, Mael Lebreton, Ruth J. van Holst, Judy Luigjes

## Abstract

Compulsivity is a common phenotype amongst various psychiatric disorders, such as obsessive-compulsive disorder (OCD) and gambling disorder (GD). Deficiencies in metacognition, such as the inability to properly estimate ones’ own performance via well-calibrated confidence judgments could contribute to pathological decision-making in these psychiatric disorders. Earlier research has indeed suggested that OCD and GD patients reside at opposite ends of the confidence spectrum, with OCD patients exhibiting underconfidence, and GD patients exhibiting overconfidence. Recently, several studies established that motivational states (e.g. monetary incentives) influence metacognition, with gain (respectively loss) prospects increasing (respectively decreasing) confidence judgments. Here, we reasoned that the OCD and GD symptomatology might correspond to an exacerbation of this interaction between metacognition and motivational states. We hypothesized GD’s overconfidence to be exaggerated during gain prospects, while OCD’s underconfidence to be worsened in loss context, which we expected to see represented in ventromedial prefrontal cortex (VMPFC) blood-oxygen-level-dependent (BOLD) activity. We tested those hypotheses in a task-based functional magnetic resonance imaging (fMRI) design. Our initial analyses showed increased confidence levels for GD versus OCD patients, that could partly be explained by sex and IQ. Although our primary analyses did not support the hypothesized interaction between incentives and groups, exploratory analyses did show increased confidence in GD patients specifically in gain context. fMRI analyses confirmed a central role for VMPFC in the processing of confidence and incentives, but with no differences between the clinical samples. The trial is registered in the Dutch Trial Register (Trial NL6171, registration number: NTR6318) (https://www.trialregister.nl/trial/6171).

## Introduction

Compulsive behaviors are defined as “repetitive acts that are characterized by the feeling that one ‘has to’ perform them while being aware that these acts are not in line with one’s overall goal”^1^. Various psychiatric disorders are associated with compulsivity, of which obsessive-compulsive disorder (OCD) is the most typical^2^, but it’s also seen in addictive disorders such as gambling disorder (GD)^3^. Both disorders are characterized by a loss of control over their compulsive behaviors, albeit originating from distinct motivations, serving different purposes and relating to distinct symptomatology^4,5^. Hence, compulsivity seems to be a common phenotype in otherwise symptomatically different disorders.

Dysfunctions in metacognition could explain distinct features of compulsive behaviors. Metacognition is the ability to monitor, reflect upon and think about our own behavior^6^. One metacognitive computation is the judgment of confidence, defined as the subjective estimate of the probability of being correct about a choice^7^. Confidence plays a key role in decision-making and learning^6–8^, and therefore in steering our future behavior^9,10^. It is crucial for behavioral control that one’s confidence is in line with reality. Nonetheless, discrepancies between actual behavior (e.g. choice accuracy) and confidence in that behavior (subjective estimate of accuracy) have been consistently described, which could contribute to pathological (compulsive) decision-making as seen in various psychiatric disorders^11^. Clinical presentations of OCD and GD indeed suggest confidence abnormalities in opposite direction, under- and overconfidence, respectively, which could both promote detrimental decision-making, such as checking behavior and compulsive gambling^12–15^. In a recent review we showed that both people with subclinical and clinical OCD consistently showed a decrease in confidence level, which was especially profound in OCD-symptom contexts^11^. Oppositely, in pathological gamblers, there was evidence for overconfidence in rewarding gambling contexts. which was also related to symptom severity^16,17^. In sum, GD and OCD patients seem to function at opposite sides of the confidence continuum, respectively over- and under-estimating their performance, which could explain how opposite traits may underlie similar pathological behavior (i.e. compulsive behavior).

Reward processes are important for learning and decision-making and interact with cognition^18^. Many studies have implicated subcortical regions such as the ventral striatum (VS) and cortical regions such as the ventromedial prefrontal cortex (VMPFC) in reward processing, forming a ‘brain valuation system’^19–21^ whose activity relates to value-based decision-making^22^ and motivates behavior^23^. Both OCD and GD patients show deficits in reward processes and accompanying dysregulated neural circuitries. A recent review on neuroimaging of reward mechanisms by Clark et al. (2019) clearly indicated dysregulated reward circuitries, especially focused on the VMPFC and VS in GD, with mixed evidence regarding the direction of these effects^24^. In OCD, a recent review showed that the ventral affective circuit, consisting of medial frontal cortex and VS was consistently shown to be dysregulated, showing decreased activity in response to rewards, which was increased in response to losses^25^. This is particularly relevant to the question of how confidence might contribute to those pathologies’ symptoms, as an increasing number of studies show that affective and motivational states can influence confidence^26–28^. Recently, we demonstrated that monetary incentives bias confidence judgments in healthy individuals, where prospects of gain (respectively loss) increase (respectively decrease) confidence, whilst performance levels remained unaffected in both perceptual and reinforcement-learning contexts^29–32^.

We therefore reasoned that an interaction between incentive and confidence processing could cause or fuel the compulsive behaviors in GD and OCD. On the one hand, prospects of high monetary incentives could exaggerate overconfidence in GD patients, leading to continuation of compulsive gambling. On the other hand, in OCD this could lead to exaggerated decreased confidence in negative value context as harm avoidance is considered one of the core motivations of compulsive behavior in OCD^33–35^.

On the neurobiological side, a growing number of functional magnetic resonance imaging (fMRI) studies have associated metacognitive processes with activity in the frontal-parietal network^36–40^, and activity in the dorsomedial prefrontal cortex (dmPFC), insula and dorsal anterior cingulate cortex (dACC) has been negatively associated to confidence, suggesting a role for these areas in representing uncertainty-related variables^41–45^. Interestingly, recent studies have also found activity in the VS, the VMPFC and perigenual anterior cingulate cortex (pgACC) - to be positively associated with confidence^41,46–51^. Importantly, this latter network has been previously positively associated with value-based processes^20,21,52,53^. Actually, both confidence judgments and value information seem to be automatically integrated into VMPFC’s activity^20,22,47,54,55^. Yet, little is known about if and how the behavioral interaction observed between incentives and confidence can be explained by their shared association with the VMPFC. In an attempt to answer this question, we recently reported an important interaction between incentive and metacognitive signals in the VMPFC in healthy subjects: confidence signals in the VMPFC were observed in trials with gain prospects, but disrupted in trials with no – or negative (loss) monetary prospects^30^. This suggest that the VMPFC has a key role in mediating the relation between incentives and metacognition. Given the crucial roles of the VMPFC and VS in reward processes and metacognition, which were found to be dysregulated in GD and OCD, we hypothesized that both regions would show disrupted activation patterns related to incentive processing and metacognition and their interaction in patients compared to healthy controls.

Overall, in the present study we investigate metacognitive ability and its interaction with incentive motivation in OCD and GD, behaviorally and neurobiologically.

## Methods

### Ethics

Experimental procedures were approved by the Medical Ethics Committee of the Academic Medical Center, University of Amsterdam. All subjects provided written informed consent.

### Participants

We recruited a total of 31 GD patients, 29 OCD patients and 55 HCs between 18 and 65 years old. Of our HC sample of 55 subjects, 25 subjects were included in our earlier work^30^. HCs were recruited through online advertisements and from our participant database. GD patients were recruited from a local treatment center (Jellinek Addiction Treatment Center Amsterdam) and were recently diagnosed with GD. OCD patients were recruited through the department of psychiatry at the Academic Medical Center in Amsterdam and were diagnosed with OCD.

### Exclusion criteria

After applying all exclusion criteria (see Supplementary Materials), we included 27 GD patients, 28 OCD patients and 55 HCs for the behavioral analyses, of which four, two and two participants contributed only one of two task sessions, respectively. For the fMRI analyses we included 24 GD patients, 27 OCD patients and 53 HCs, of which seven, three and two participants contributed only one of two task sessions, respectively.

### Experimental Design and Study Procedure

We used a similar experimental design and study procedure as previously described^30^. For details on the experimental design and study procedure, see Hoven et al. (2020) and **Figure 1**. In sum, subjects performed a simple perceptual decision-making task, with a 2-alternative forced choice of contrast discrimination followed by a confidence judgment. In each trial, participants could either win (gain context) or lose (loss context) points, or not (neutral context), conditional on the accuracy of the choice in that trial. Importantly, this incentivization was administered after the choice moment, but before the confidence rating. The task was implemented using MATLAB® (MathWorks Inc., Sherborn, MA, USA) and the COGENT toolbox.

**Figure 1.**
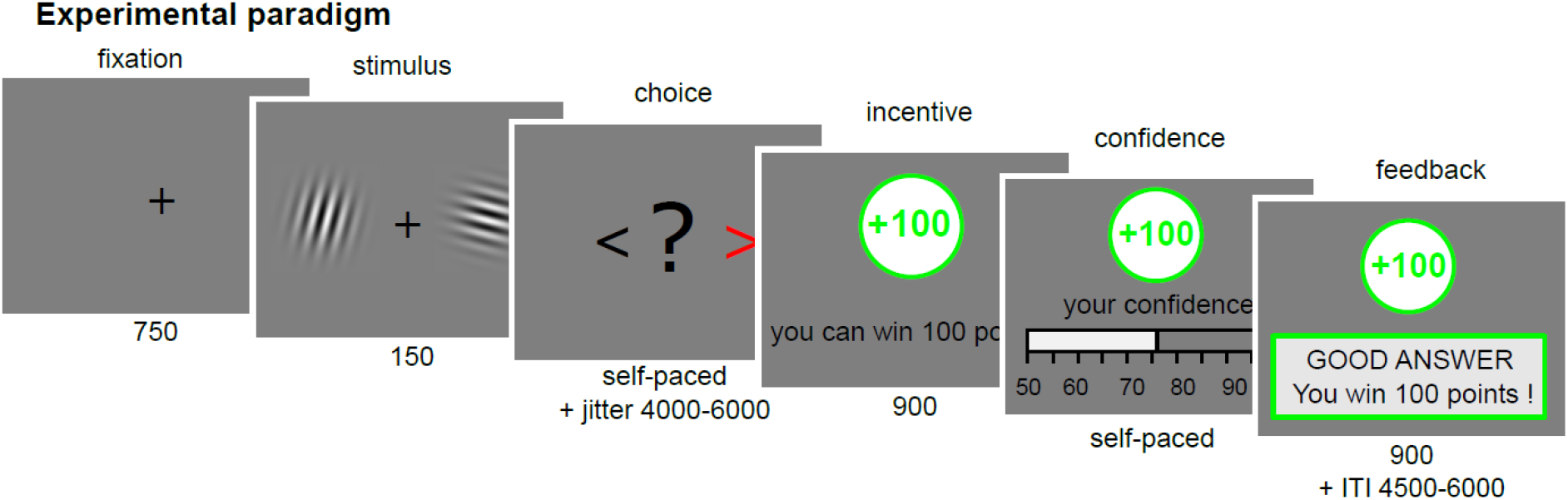
Experimental paradigm. Participants viewed two Gabor patches on both sides of the screen (150 ms) and then chose which had the highest contrast (left/right, self-paced) (for more information, see Hoven et al., 2020). After a jitter of a random interval between 4500 to 6000 ms, the incentive was shown (900 ms; green frame for win trials, grey frame for neutral trials, red frame for loss trials). Afterwards, participants were asked to report their confidence in their choice on a rating scale ranging from 50% to 100% with steps op 5%. The initial position of the cursor was randomized between 65% and 85%. Finally, subjects received feedback. The inter trial interval (ITI) had a random duration between 4500 and 6000 ms. The calibration session only consisted of Gabor discrimination, without confidence rating, incentives or feedback and was used to adjust difficulty so that every individual reached a performance of 70%.

### Behavioral Measures

We extracted trial-by-trial experimental factors: incentive condition, evidence and behavioral measures: accuracy, confidence ratings, reaction times. Evidence was calculated by normalizing the unsigned difference of the two Gabor patches’ contrast intensities by their sum to adjust for saturation effects (for more details see^31^). In addition, we computed an extra *latent* variable: early certainty.

The early certainty variable was computed in order to analyze BOLD activity at choice moment, when the brain encodes a confidence signal that is not yet biased by incentives. This was done by making a trial-by-trial prediction of early certainty based on stimulus features (reaction times, evidence and accuracy) at choice moment. This resulted in an early certainty signal that was highly correlated with confidence, but showed no statistical relationship with incentives (see Supplementary Materials). For more details, see^30^.

Next to confidence ratings we also assessed additional metacognitive metrics:(1) Confidence calibration, the difference between average confidence and average performance as an indicator of over- or underconfidence, (2) Metacognitive sensitivity, the ability to discriminate between correct answers and errors using confidence judgments (see Supplementary Materials).

### Behavioral Analyses

All analyses were performed in the R environment (RStudio Team (2015). RStudio: Integrated Development for R. RStudio, Inc., Boston, MA). We used linear mixed effects models (LMEMs) as implemented in the lmer function from the lme4 and afex packages^56,57^. To determine p-values for the fixed effects, we performed Type 3 F tests with Satterthwaite approximation for degrees of freedom as implemented in the afex package. When relevant, we used the ‘emmeans’ package to perform post-hoc tests that were corrected for multiple comparisons using Tukey’s method^58^.

To answer our main research questions, we built several LMEMs and performed a model selection procedure (see Supplementary Materials). The final model (Model 1) included fixed effects of incentive, group, accuracy and evidence (z-scored) and interactions between incentive and group, as well as two-way and three-way interactions between evidence, accuracy and group. Moreover, a random subject intercept and a random slope of incentives per subject were included in the final model as well. To confirm that the incentive condition or group did not influence accuracy or reaction time, we modelled additional LMEMs with performance and reaction time as dependent variables (Model 2, Model 3).

Lastly, we added IQ (z-scored) and sex as fixed effects to our original Model 1 (Model 4) to control for differences in the distribution of these demographic variables. Model fit was assessed and compared using Chi-square tests on log-likelihood values. Additional control analyses on the properties of confidence, early certainty, confidence calibration and metacognitive sensitivity are reported in the Supplementary Materials.

Due to a technical bug, our design was not fully balanced as the level of perceptual evidence was not equal across the incentive conditions. ANOVA and post-hoc testing indeed showed that evidence was highest in neutral condition, followed by gain and loss. There were no group differences, nor an interaction between group and incentive. These effects cannot account for any group differences we find in our data, since evidence did not differ between groups. Importantly, the evidence differences did not affect performance, since performance is equal across conditions. See Supplementary Materials for more details.

### fMRI analyses

For details on fMRI acquisition and preprocessing see Supplementary Materials and Hoven et al (2020)^30^.

All fMRI analyses were conducted using SPM12. Critically, our design allowed us to distinguish between our two timepoints of interest: 1) the moment of stimulus presentation and choice in which implicit (un)certainty about the choice is formed, and 2) the moment of incentive presentation and confidence rating, in which the value of incentives and the confidence rating are encoded. We built a general linear model (GLM 1) estimated on subject-level with these two moments of interest: the moment of choice (i.e. stimulus presentation) and the moment of incentive presentation/confidence rating. We chose to analyze the incentive presentation and confidence rating as a single timepoint since the rating moment followed the presentation of the incentive after 900 ms, with regressors time-locked to the onset of incentive presentation. We also included a regressor for the moment of feedback to explain variance in neural responses related to feedback on accuracy and value that was not related to the decision-making process, but this regressor was not of interest for the current analyses. All whole-brain activation maps were thresholded using family-wise error correction (FWE) at cluster level (PFWE_clu < 0.05), with a voxel cluster-defining threshold of p<.001 uncorrected.

Using GLM 1, with regressors for choice modulated by early certainty, for incentive/rating modulated by incentive and confidence, and for feedback modulated by accuracy we were able to investigate our contrasts of interest: (1) choice moment modulated by early certainty, (2) incentive/rating moment modulated by incentive value and (3) incentive/rating moment modulated by confidence rating. For details see Supplementary Materials.

In order to study the interaction between incentive motivation and metacognitive ability on the neurobiological level we leveraged the factorial design of our task to build GLM 2. We used GLM 2 to explicate the effect of incentive motivation on both the integration of evidence at choice moment, as well as on confidence formation, and compare those between groups. GLM 2 consisted of regressors for each time point (choice and incentive/rating moments) and for each incentive condition, as well as a single regressor at feedback moment, resulting in seven regressors. For all these events we examined both baseline activity and regression slopes relating to their pmod of interest: signed evidence for choice and confidence for incentive/rating. See Supplementary Materials for more details.

Since the results by Hoven et al., 2020 suggested that the VMPFC plays an important role in the interaction between incentive motivation and metacognition, we created a functional region of interest (ROI) that represented the confidence-related activity in the VMPFC cluster from our GLM 1 across groups results (see **Figure 4C, Table 5**). We then extracted individual *t*-statistics within this ROI (i.e. normalized beta estimates^59^) from our contrasts of interest and performed one-sample t-tests against 0 to check for positive or negative activation patterns. Then, we compared them between incentive conditions, groups, and studied their interactions using mixed ANOVAs implemented in the afex package. When appropriate, we performed post-hoc testing using the emmeans package, correcting for multiple comparisons using Tukey’s method. Since we also hypothesized that the VS would play a role in the interaction between incentives and metacognition, we performed the same ROI analysis in the VS with a functional ROI that represented the incentive-related activity in the VS cluster from our GLM 1 across group results (see **Table 5**).

## Results

### Demographics

IQ and sex distributions differed between groups (IQ: F_2,107_ = 3.222, p=0.0438; sex: X = 14.483, df = 2, p<.001), with higher IQ scores for HC subjects compared with GD patients (t = 2.53, p=0.014) and with mostly men in the GD group, and relatively more women in the OCD group (Table 2). This corresponds to the natural distribution observed in epidemiological studies for OCD and GD, showing higher prevalence of GD amongst men, and a slightly higher prevalence of OCD in women^60–63^. Age did not differ between groups. For post-hoc group differences on questionnaire scores, see Supplementary Materials.

**Table 1.**
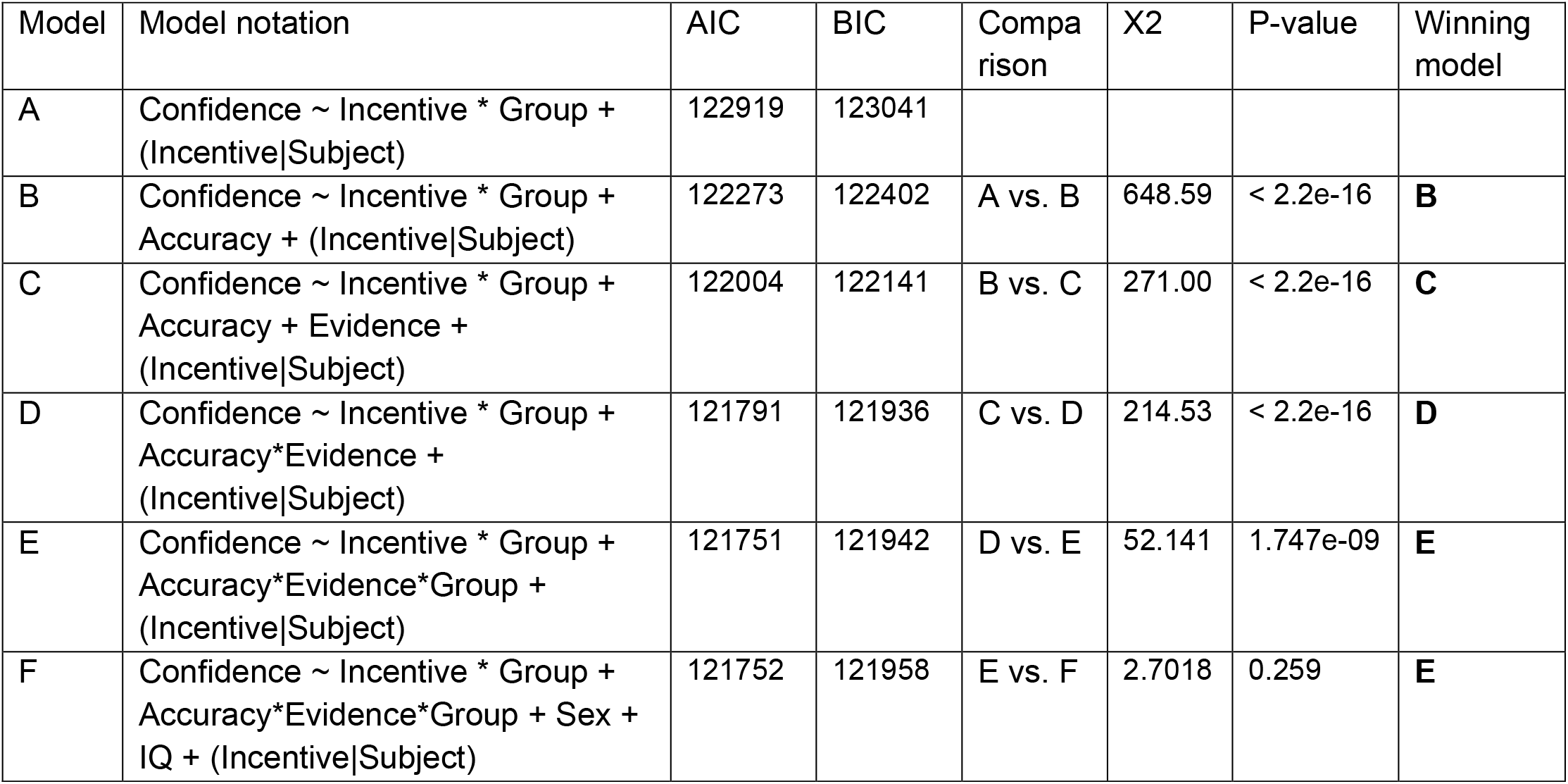
Model descriptions and comparison. Shown here are the model notations of all models with their respective Akaike Information Criterion (AIC) and Bayesian Information Criterion (BIC) values, as well as model comparison outcomes with corresponding χ2 and P-values, resulting in the winning model ‘E’, which is referred to as Model 1 in the manuscript.

**Table 2.**
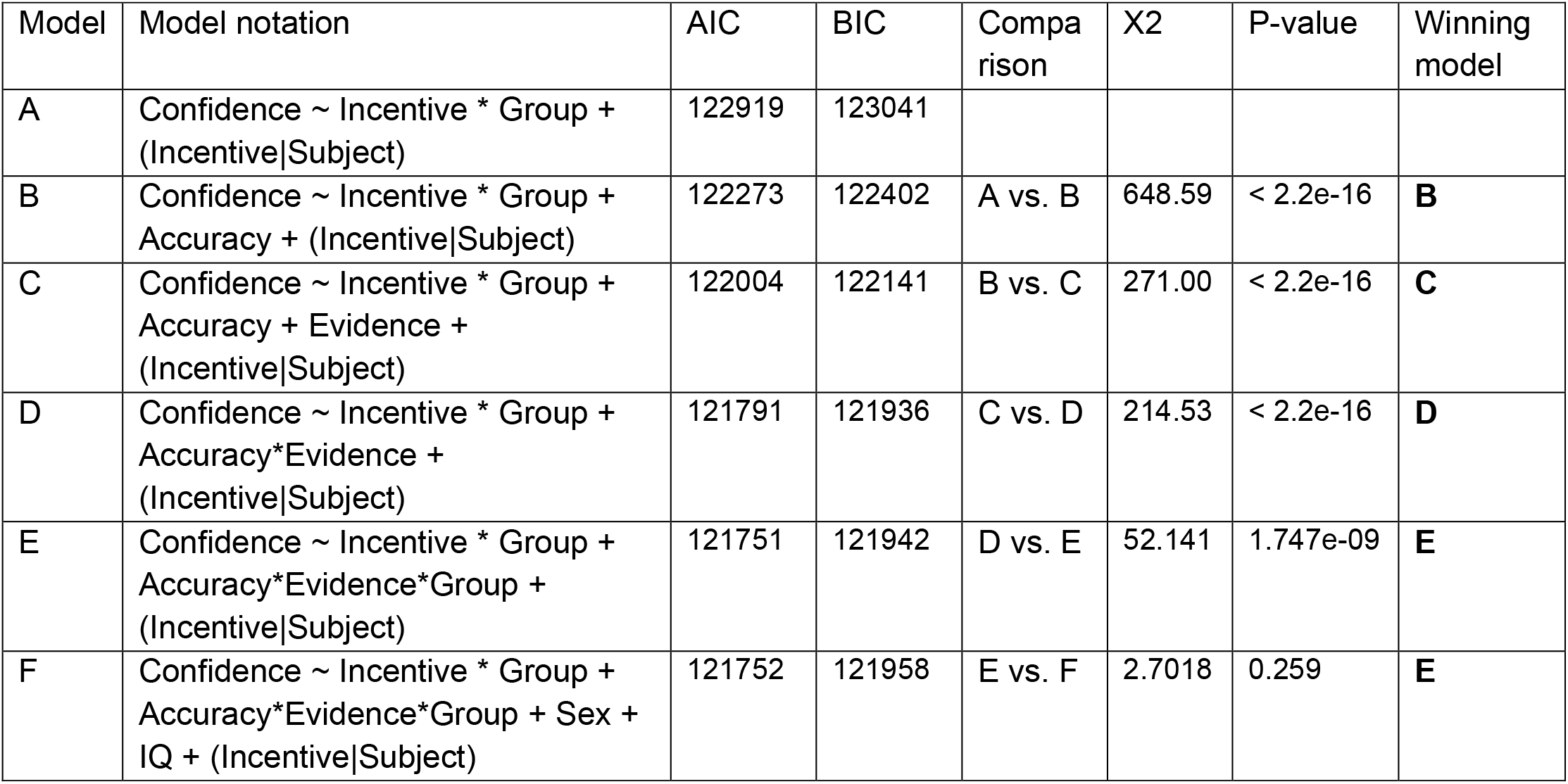
Demographics: Means +-standard deviations of various demographic variables are shown per group, for sex counts are displayed. Statistics for group comparisons are shown, including F and X^2^ statistics, degrees of freedom and p-values. IQ= estimated Intelligence Quotient, GD = gambling disorder, HAMA = Hamilton Anxiety Rating Scale, HC = healthy control, HDRS = Hamilton Depression Rating Scale, OCD = obsessive-compulsive disorder PGSI = Problem Gamblers Severity Index, Y-BOCS = Yale-Brown Obsessive Compulsive Scale. *p<.05, ***p<.001

### Behavioral Results

To start, we answered our main questions: (1) are there group differences in confidence, and (2) what is the influence of incentive motivation on confidence. Model 1 showed a main effect of group (F_2,112_ = 4.7910, p=.01) and incentive (F_2,112_ = 20.9371, p<.001) on confidence (**Figure 2, Supplementary Table 3**). We also found a main effect of accuracy (F_1,15107_ = 608.8906, p<0.001), with subjects showing higher confidence for correct answers. Moreover, there was a significant two-way interaction of group and evidence (F_2,15099_ = 3.5094, p=0.02994). As expected, we also found a significant interaction between accuracy and evidence, replicating the ‘X-pattern’ signature of evidence integration where confidence increases with increasing evidence when correct, and vice versa (F_1,15097_=185.3245, p<0.001)^64^. Interestingly, the evidence integration effect differed per group, as signaled by a significant three-way interaction between accuracy, evidence and group (F_2,15094_ = 3.0533, p=0.04723) (**Supplementary Figure 3, Supplementary Table 3**, for post-hoc tests see Supplementary Materials). Lastly, the interaction between incentive and group revealed a trend towards an effect (F_4,112_= 2.2821, p=0.06487).

**Figure 2.**
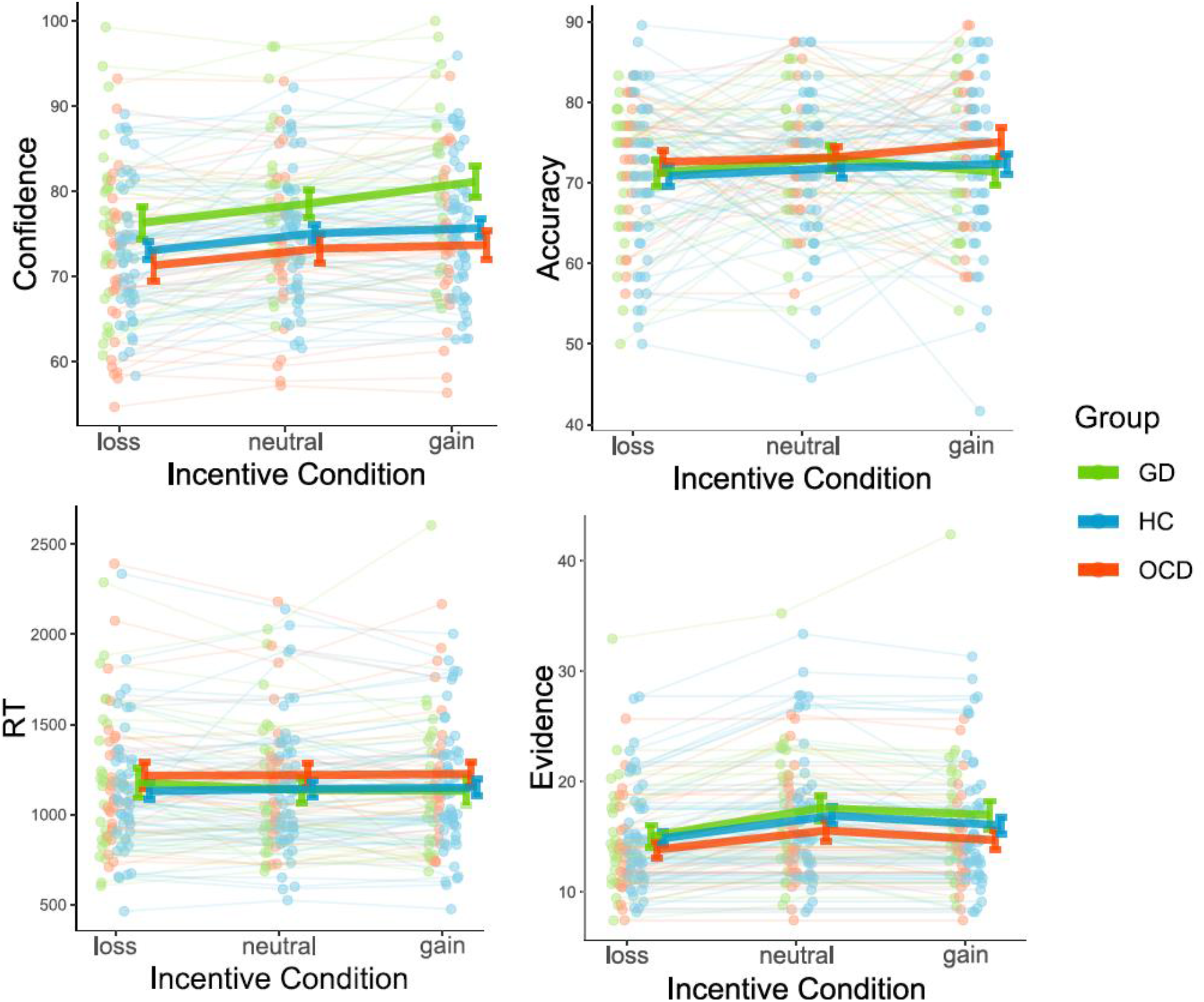
Behavioral results. Individual-averaged confidence, accuracy, reaction times and evidence as a function of incentive condition (loss, neutral and gain) per group. Green dots and lines represent gambling disorder patients, blue dots and lines represent healthy controls and red dots and lines represent obsessive-compulsive disorder patients. Dots represent individuals, and lines highlight within subject variation across conditions. Error bars represent sample mean ± SEM per group. GD = gambling disorder, HC = healthy control, OCD = obsessive-compulsive disorder

Post-hoc tests indicated a significantly higher confidence in GD patients versus OCD patients (GD-OCD = 6.38 +-2.12, Z-ratio = 3.014, p=0.0073), and a trend towards higher confidence in GD compared to HC subjects (GD-HC = 4.30 +- 1.84, Z-ratio = 2.333, p=0.0513), whereas OCD patients did not differ from HC subjects. Moreover, we replicated the parametric effect of incentive value on confidence (loss-neutral =-1.80 +- 0.429, Z-ratio = -4.192, p<0.001; loss-gain =-3.14 +- 0.486, Z-ratio = -6.460, p<0.001; neutral-gain = -1.34 +- 0.363, Z-ratio = - 3.683, p<0.001). With regards to the three way interaction we found that GD patients’ confidence was less influenced by evidence for correct answers compared to both HCs and OCD patients (see Supplementary Materials, **Supplementary Figure 3**). Exploratory post-hoc analyses on the group*incentive interaction effect showed that, especially in context of possible gains, GD patients were more confident than OCD patients (GD - OCD = 8.12 +-2.24, Z-ratio = 3.621, p<0.001) and HC subjects (GD - HC = 5.83 +- 1.95, Z-ratio = 2.989, p=0.0079), with no differences between HC and OCD patients in any incentive condition (**Table 3**).

**Table 3.** Results of linear mixed-effects models. Shown here are the results of Model 1 (without demographics) and Model 4 (with demographics) acquired using Type 3 F tests with Satterthwaite approximation for degrees of freedom using the afex package. Shown are F values, with corresponding degrees of freedom and P-values.

| **Model 1** | Confidence |
| --- | --- |
| Incentive | $F(2.00, 112.34) = 20.94, p < .001$ |
| Group | $F(2.00, 112.51) = 4.79, p = .010$ |
| Accuracy | $F(1.00, 15107.05) = 608.89, p < .001$ |
| Evidence | $F(1.00, 15104.05) = 0.04, p = .848$ |
| Incentive:Group | $F(4.00, 112.10) = 2.28, p = .065$ |
| Accuracy:Evidence | $F(1.00, 15097.33) = 185.32, p < .001$ |
| Group:Accuracy | $F(2.00, 15106.28) = 2.27, p = .103$ |
| Group:Evidence | $F(2.00, 15099.41) = 3.51, p = .030$ |
| Group:Accuracy:Evidence. | $F(2.00, 15094.35) = 3.05, p = .047$ |
| **Model 4** | Confidence |
| Incentive | $F(2.00, 112.34) = 20.93, p < .001$ |
| Group | $F(2.00, 112.50) = 2.75, p = .068$ |
| Sex | $F(1.00, 110.26) = 2.88, p = .093$ |
| IQ | $F(1.00, 109.80) = 0.03, p = .865$ |
| Accuracy | $F(1.00, 15107.01) = 609.14, p < .001$ |
| Evidence | $F(1.00, 15104.51) = 0.04, p = .845$ |
| Incentive:Group | $F(4.00, 112.11) = 2.29, p = .064$ |
| Accuracy:Evidence | $F(1.00, 15097.16) = 185.42, p < .001$ |
| Group:Accuracy | $F(2.00, 15106.06) = 2.30, p = .100$ |
| Group:Evidence | $F(2.00, 15098.91) = 3.45, p = .032$ |
| Group:Accuracy:Evidence | $F(2.00, 15094.15) = 3.09, p = .046$ |

As control analyses we estimated Model 2 and 3 with accuracy and reaction time as dependent variables.(**Table 4**). No effect of group, incentive or an interaction effect on accuracy or reaction time were found, as expected from our design (where incentives follow choices), confirming that accuracy and response times cannot confound any effect of incentives that we found on confidence.

**Table 4.** Results of control models. Shown here are the results of Model 2 and Model 3 linear mixed-effects models, acquired using Type 3 F tests with Satterthwaite approximation for degrees of freedom using the afex package. Shown are F values, with corresponding degrees of freedom and P-values

|  |  |
| --- | --- |
| **Group** | $F_{2,109} = 0.5827, P = 0.5601$ |
| **Incentive** | $F_{2,1591} = 1.0319, P = 0.3566$ |
| **Group\*Incentive** | $F_{4,1586} = 0.8671, P = 0.4830$ |

|  |  |
| --- | --- |
| **Group** | $F_{2,110} = 0.5207, P = 0.5956$ |
| **Incentive** | $F_{2,220} = 0.0994, P = 0.9054$ |
| **Group\*Incentive** | $F_{4,219} = 0.4269, P = 0.7891$ |

Since sex and IQ were significantly different between the groups, we aimed to control for these variables by adding them as fixed effects, resulting in Model 4. The main effect of group did not remain significant, but showed a trend towards an effect (F2,112 = 2.7465, p=0.06846), while the main effect of incentive did remain significant (F2,112 = 20.9326, p< 0.001). We found no evidence for a significant effect of sex (F1,110 = 2.8776, p=0.09264), or IQ (F1,109 = 0.0291, p=0.86489). The interaction effect between group and incentive remained non-significant at trend-level (F4,112 = 2.2898, p=0.06412). The significant three-way interaction between accuracy, evidence and group persisted (F2,15094 = 3.0871, p=0.04566). Importantly, when performing a Chi-square test on the log-likelihood values of the models excluding and including the demographic variables to compare model fit, the model without demographics showed a better model fit (X2 = 2.7018, df=2, p=0.259), thereby favoring this simpler model. Additionally, to investigate how confidence was differently affected by sex in our healthy controls, we performed a two-sample t-test which showed that males were generally more confident than females (males: 76.51 +- 1.04; females: 71.70 +- 0.77) (t_52_ = 2.6518, p-value=0.01057). However, both sex and IQ did not show a significant influence on confidence level in Model 4.

Next to confidence, we also examined calibration and metacognitive sensitivity (see Supplementary Materials). In short, we showed that GD patients were more overconfident than OCD patients, without an effect of incentive condition. No differences in metacognitive sensitivity were found between groups or incentive conditions.

### fMRI results GLM 1

We analyzed functional neuroimaging data to test for differences in brain activity between groups for our contrasts of interest: (1) choice moment modulated by early certainty, (2) rating/incentive moment modulated by incentive value, and (3) rating/incentive moment modulated by confidence. The results from the fMRI group analysis revealed no significant differences between the groups for any of our contrasts.

Next, we grouped all subjects together and performed one-sample t-tests on our contrasts of interest to examine the results across groups (cluster-generating voxel threshold p<.001 uncorr.; clusterwise correction for multiple comparisons p_FWE_<0.05). During choice, early certainty positively correlated with activation in the precuneus, VMPFC, bilateral VS and putamen, and bilateral visual areas (**Figure 3A**). The dorsal anterior cingulate cortex, bilateral dorsomedial- and dorsolateral prefrontal cortex, bilateral insula, thalamus, middle frontal gyrus, bilateral sensorimotor cortex, superior and inferior parietal lobe related negatively to early certainty (**Figure 3A**).

**Figure 3.**
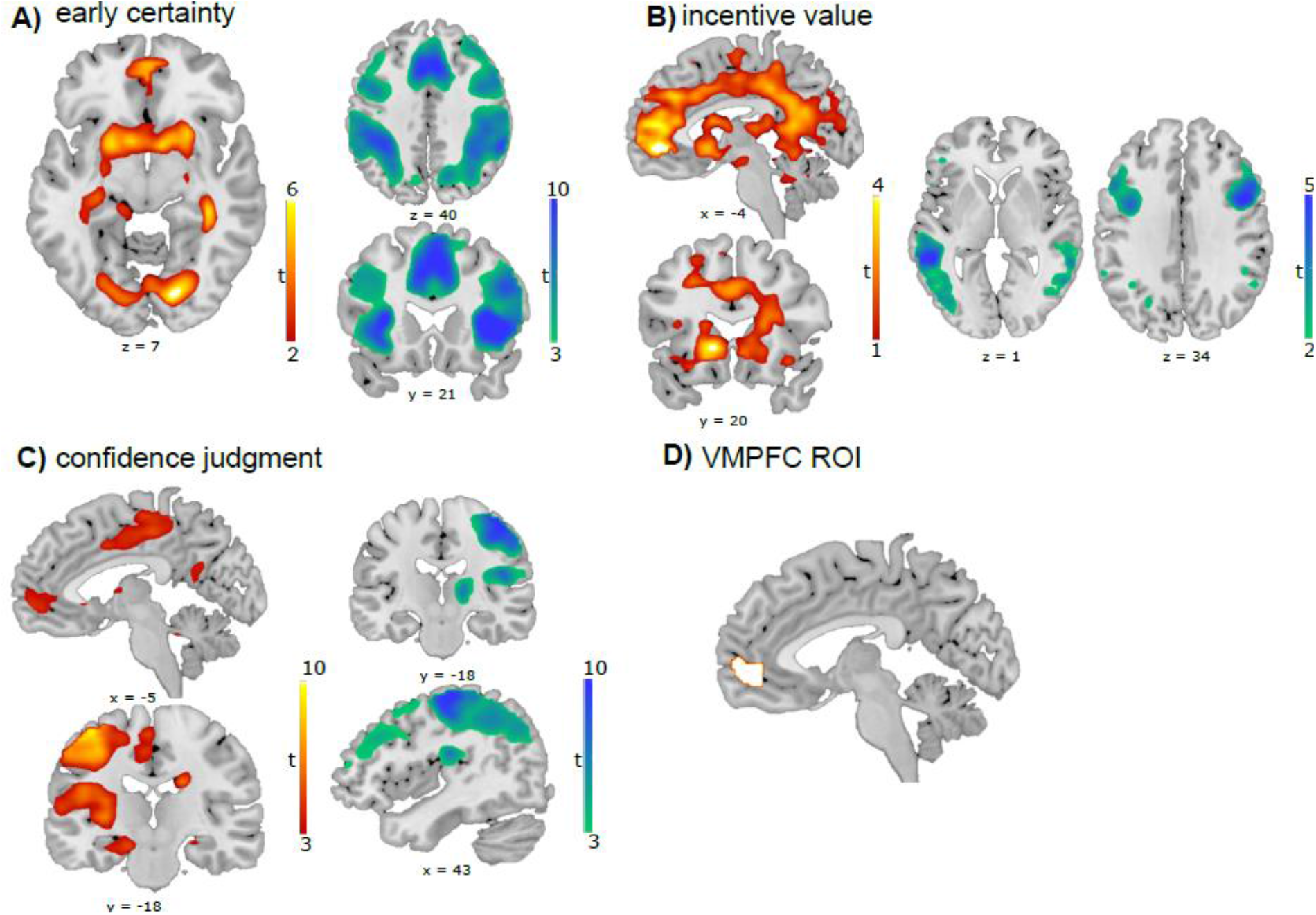
Whole brain statistical bold-oxygen-level-dependent (BOLD) activity across groups. Red/yellow areas represent areas with a positive relationship, while green/blue areas represent areas that have a negative relationship. (A) Areas correlating significantly with early certainty at choice moment. Shown are positive activations in ventromedial prefrontal cortex, ventral striatum and visual cortices. Negative activations in dorsal anterior cingulate cortex, dorsolateral prefrontal cortices, insula, parietal cortices. (B) Areas correlating significantly with incentive value at incentive/rating moment. Shown are positive activations in ventromedial prefrontal cortex, anterior cingulate cortex, ventral striatum. Negative activations in dorsolateral prefrontal cortices and temporal gyri (C) Areas correlating significantly with confidence judgments at incentive/rating moment. Positive actions are shown in ventromedial prefrontal cortex, motor cortex and putamen. Negative clusters in motor cortex and dorsolateral prefrontal cortex. All clusters survived P<0.05 FWE cluster correction. Voxel-wise cluster-defining threshold was set at P<.001, uncorrected. For whole brain activation table see table 5. (D) Region of interest (ROI) of the VMPFC used for GLM2 analyses.

At the moment of incentive presentation, the incentive value correlated positively with activation in the VS and VMPFC stretching into more dorsal areas, as well as the superior temporal gyrus (**Figure 3B**). Incentive value was negatively related to activity in the right (pre)motor cortex and dorsolateral PFC, as well as the left middle and superior temporal gyrus, left occipitotemporal gyrus, and left middle and inferior frontal gyrus. Moreover, activity in right lateral occipitotemporal gyrus and middle temporal gyrus were negatively related to incentive value (**Figure 3B**).

During rating moment, confidence was positively related to activity in the VMPFC, left motor cortex and putamen and bilateral visual areas (**Figure 3C**). The following areas were negatively related to confidence: the left superior and inferior parietal lobes, right dorsolateral PFC, right supramarginal gyrus and thalamus, right motor cortex stretching into the dorsolateral PFC, left visual cortex and cerebellum (**Figure 3C**). See **Table 5** for details of across group fMRI results.

**Table 5.**
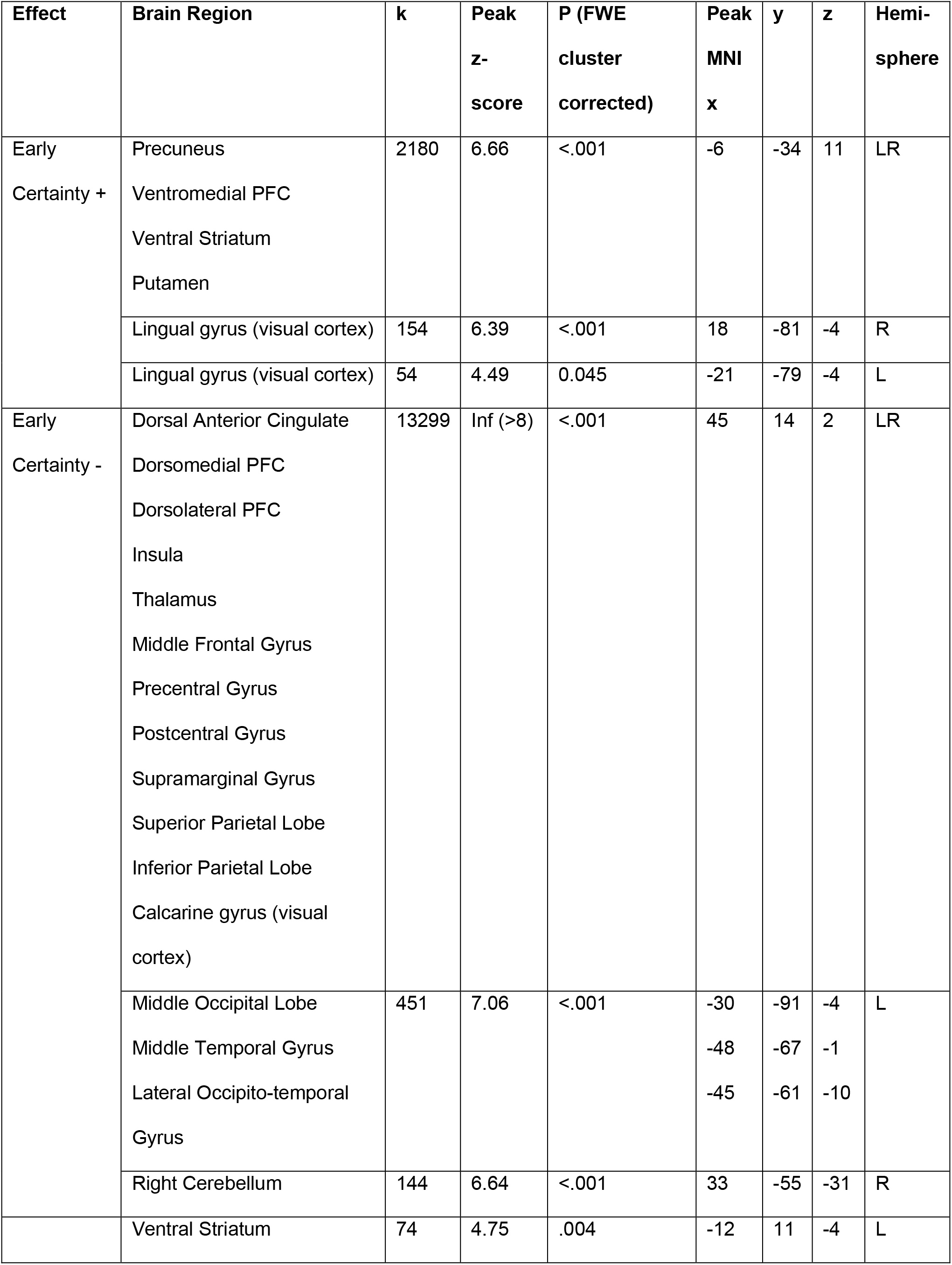
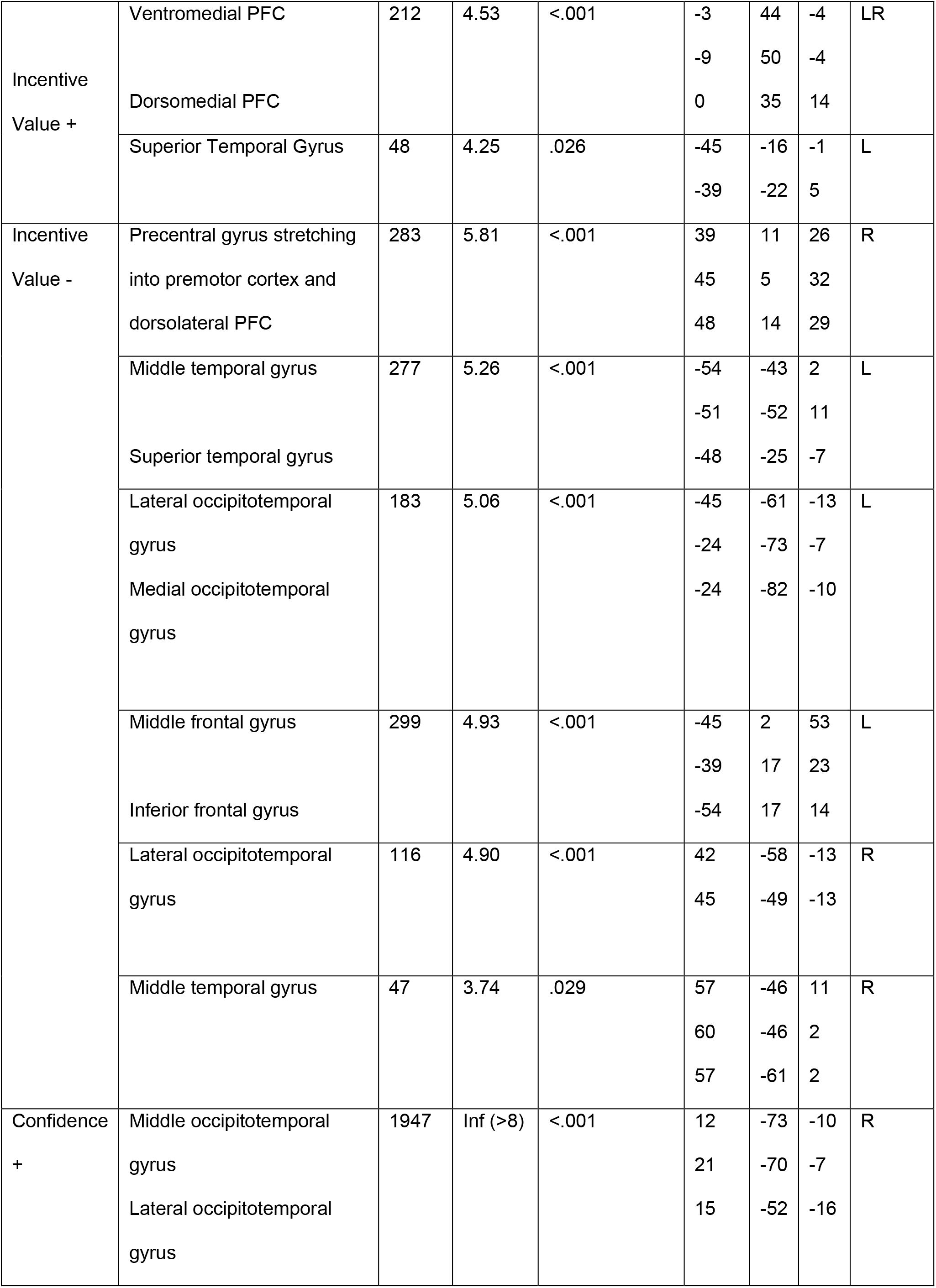
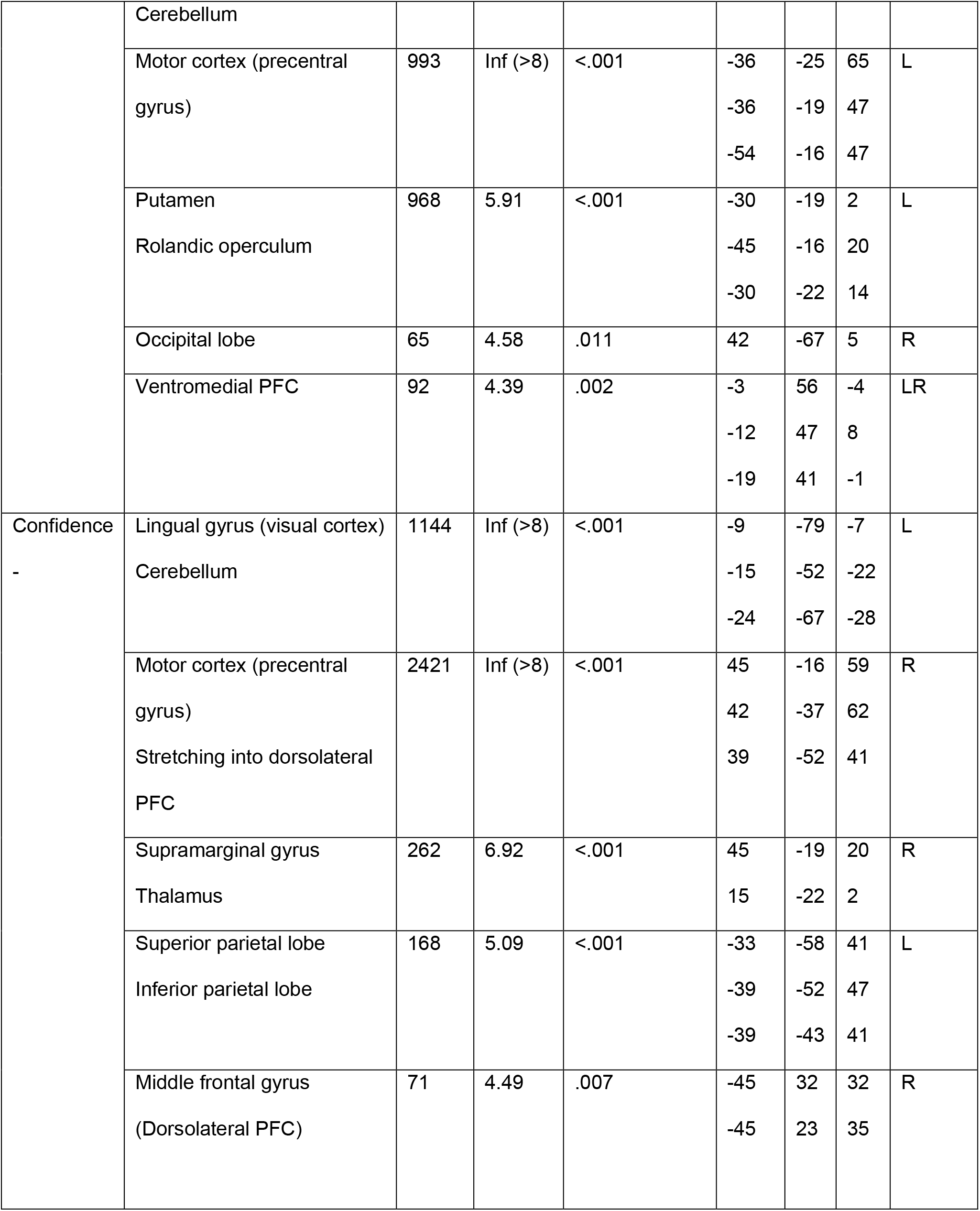
Whole brain activation tables. Brain activations (whole brain analyses) showing activity related to early certainty at choice moment, as well as activity related to incentive and confidence at incentive/rating moment. All whole-brain activation maps were thresholded using family-wise error correction for multiple correction (FWE) at cluster level (P FWE_clu < 0.05), with a voxel cluster-defining threshold of P<0.001 uncorrected. Activity that positively correlates to given variable is denoted by ‘+’, whereas negative correlations are denoted by ‘-’. PFC = prefrontal cortex.

### Interaction between metacognition and incentives in VMPFC (GLM 2)

Our recent study suggested an important role of the VMPFC in the interaction between incentive-processing and metacognitive signals^30^. To investigate how this interaction takes effect in and differs between our clinical groups, we performed an ROI analysis by leveraging our factorial design. We extracted VMPFC activations for both time points (choice and rating), all incentives (loss, neutral and gain), and all groups (HC, OCD and GD), for both baseline activity and a regression slope with (1) signed evidence and (2) confidence judgments (see **Figure 3D** for the ROI).

First, one-sample t-tests showed that, overall, VMPFC baseline activations were negative at choice and rating moment (choice: t_100_ = -3.611, p<0.001; baseline: t_100_ = -4.9287, p<0.001). The correlations between VMPFC activity and both signed evidence at choice moment and confidence at rating moment, however, were significantly positive (choice: t_100_ = 3.057, p=0.003; baseline: t_100_ = 3.7399, p<0.001) (**Figure 4**). This implies that the VMPFC represents both confidence judgments and signed evidence (i.e. interaction between accuracy and evidence: increased VMPFC activity with increased evidence when correct and vice versa).

**Figure 4.**
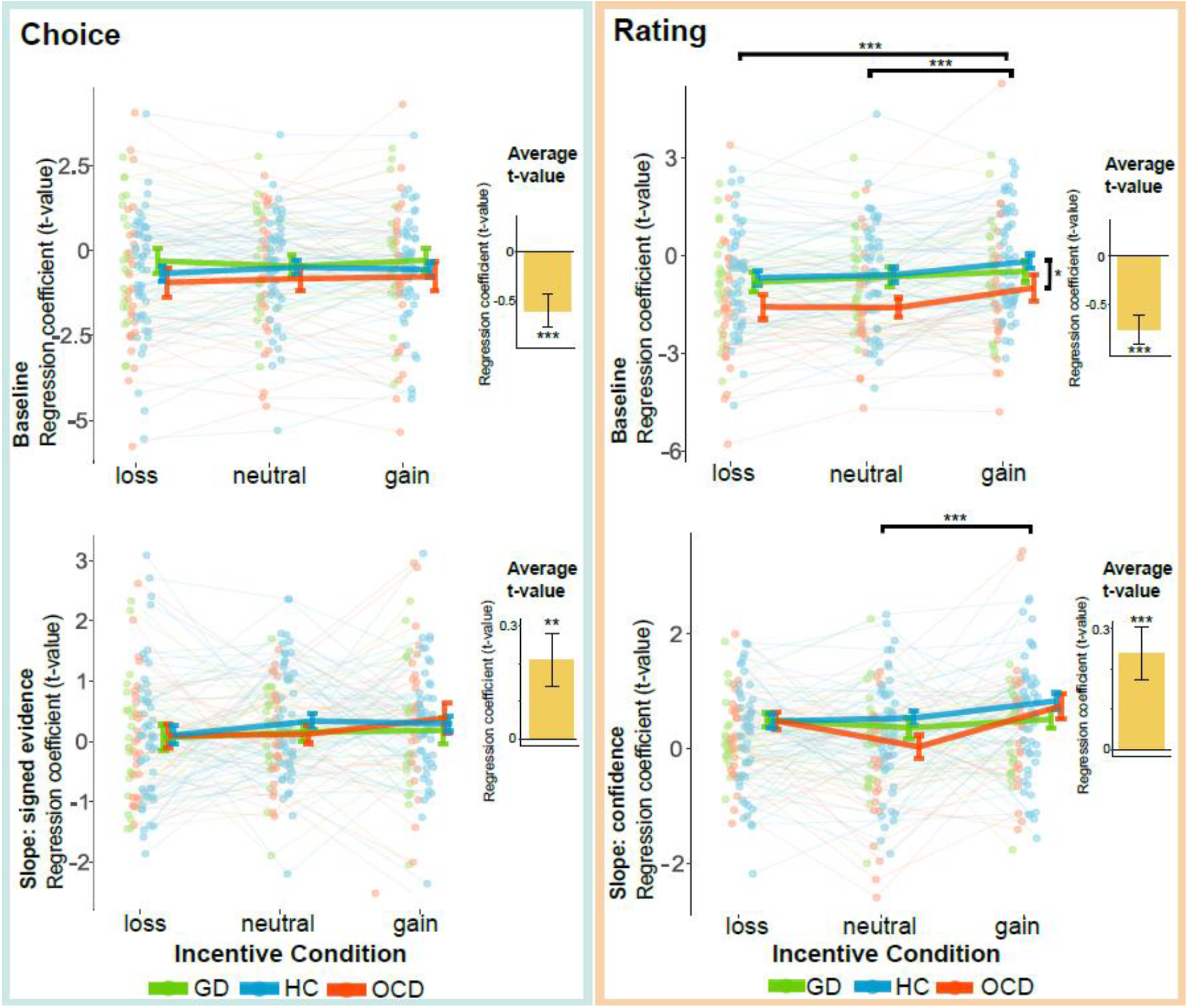
Ventromedial prefrontal cortex region of interest (ROI) analysis. T-values corresponding to baseline and regression slopes were extracted for all three groups and three incentive conditions, at two time points of interest: choice and incentive/rating moment. Green dots and lines represent gambling disorder patients, blue dots and lines represent healthy controls and red dots and lines represent obsessive-compulsive disorder patients. Dots represent individual t-statistics, and error bars represent sample mean ± SEM per group. Black bars represent significant post-hoc tests. Yellow bars represent average t-values, with corresponding significance level of one-sample t-tests against 0. (* p<0.05, ** p<0.01, *** p<0.001). GD = gambling disorder, HC = healthy control, OCD = obsessive-compulsive disorder.

Then, we investigated whether there were effects of incentive condition and group around this general signal. As expected, at choice moment there were no effects of incentive condition on VMPFC baseline activity, nor on its correlation with the signed evidence signal (i.e. slope) (**Figure 4, Table 6**). Despite the behavioral group effect on evidence integration, we did not find a group nor interaction effect on both baseline VMPFC activity and the correlation with signed evidence. At rating moment, however, incentive condition had a significant effect on both the baseline VMPFC activity, as well as its correlation with confidence. Post-hoc testing showed that the baseline VMPFC activity was higher during gain versus loss (t_196_: -3.874, p<0.001), and during gain versus neutral (t_196_ = -3.228, p<0.001), but no differences between neutral and loss conditions were found (t_196 =_ -0.646, p=0.7948). The correlation of VMPFC activity with confidence was significantly higher (i.e. increased slope) in gain versus neutral (t_196_ = -3.053, p=0.0072), while no differences between gain and loss, or between neutral and loss were found. Moreover, there was a significant group effect on VMPFC baseline activity during rating moment. The post-hoc tests revealed that OCD subjects had significantly decreased activity compared with HCs, averaged over incentive conditions (t_98_ = -2.515, p=0.0358). No interaction effects between group and incentive were found on baseline activity or its correlation with confidence at rating moment.

**Table 6:**
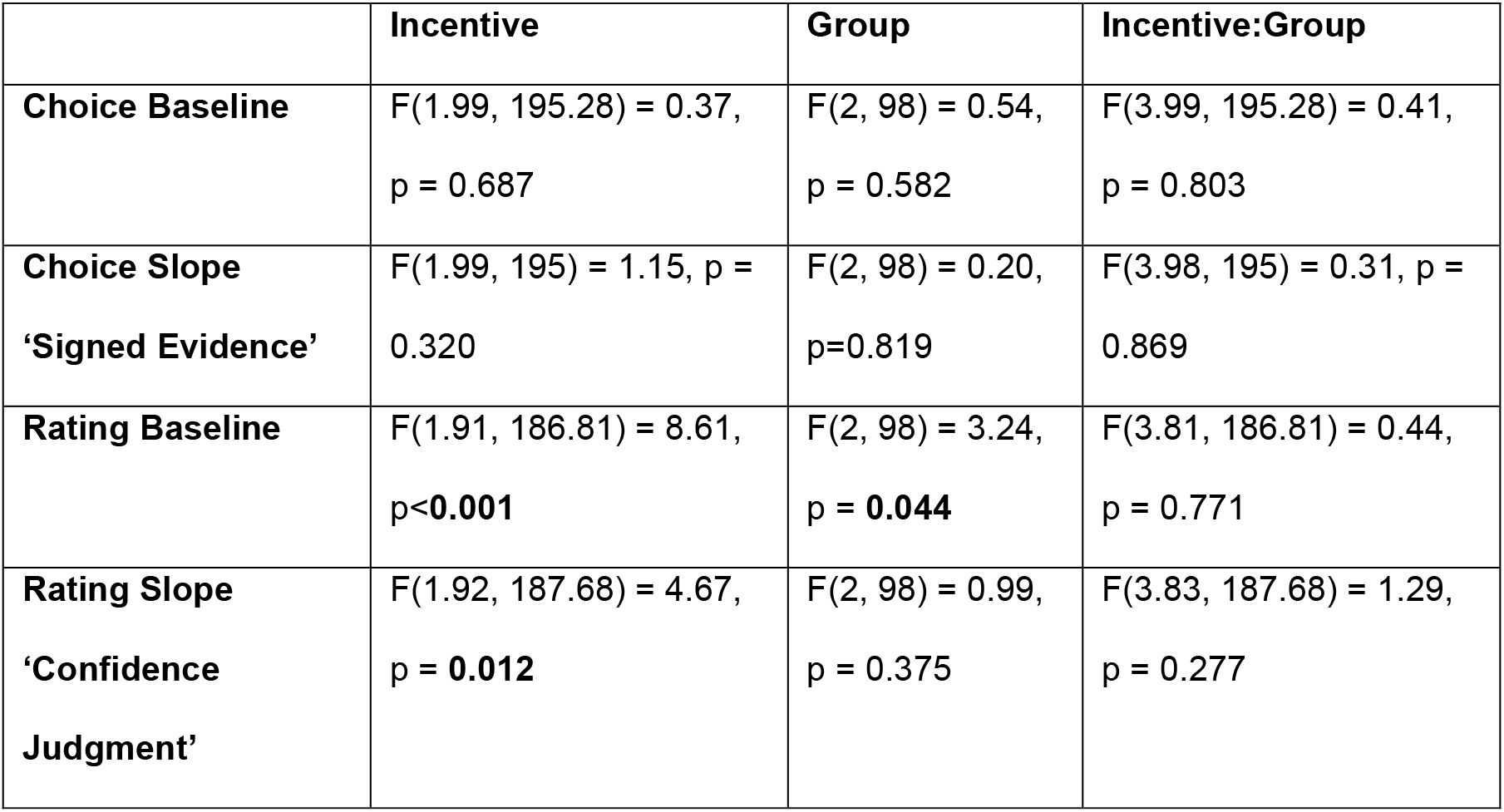
Results of VMPFC ROI analysis. Shown here are the results of the mixed ANOVAs of t-statistics in the ventromedial prefrontal cortex (VMPFC) region of interest (ROI) using the afex package. Shown are the main effects of incentive condition, group and their interaction effect on the choice and rating time points, focusing on both the baseline activity as well as the slope of signed evidence and confidence judgments, respectively. F-values, with corresponding degrees of freedom and p-values are reported.

Similar analyses using a ROI of the VS were performed (see Supplementary Materials), with similar results: VS activity correlated with signed evidence, but no incentive, group or interaction effects were found at choice moment. Similarly, the correlation of VS activity with confidence was significantly higher in gain versus neutral, with no group difference at rating moment.

## Discussion

In this study we investigated the (neural signatures of) metacognitive ability and its interaction with incentive motivation in two compulsive disorders: OCD and GD. First, we replicated the biasing effect of incentives on confidence estimation in all groups, showing that confidence was higher in the gain context and lower in the loss context. This is a robust effect, that has now been independently replicated multiple times^29–32^. We initially found evidence for a significantly higher confidence in GD patients versus OCD patients, although this effect diminished after controlling for sex and IQ differences between groups. Hence, we only found moderate evidence for our hypothesis of group differences in confidence, as well as for our hypothesis that incentive motivation would affect confidence judgments differently in the groups. Future research should address the role of the demographic confounding factors more specifically.

When looking into the computational signatures of confidence formation in more detail, GD patients interestingly showed less integration of evidence into their confidence judgments for correct choices compared to both HCs and OCD patients. This suggests that GD patients were less able to use evidence they received to form confidence judgments. This decreased sensitivity to objective evidence could fit GD’s symptomatology of cognitive inflexibility^3,65^, and cognitive distortions^66,67^. Illusion of control leads pathological gamblers to believe they can predict outcomes, rendering them less influenced by objective evidence, which may promote continuation of (overconfident) gambling behavior^13,68^.

Notably, our patient groups seemed to be situated on opposite sides of the confidence spectrum, with GD patients being more confident than OCD patients. However, this effect was partly driven by sex and IQ differences between groups. The GD group consisted mostly of males, whereas the OCD group had a more mixed composition. mirroring the prevalence distribution of these disorders^69–72^. Consistent with our findings of increased confidence in HC male subjects, recent studies have shown that males are more confident than females, despite equal performance^74,75^. Therefore, the effect of sex might have explained some variance in our data, but does not fully explain the group differences, since we do find a trend toward a group effect. The importance of taking into account sex and gender as factors in both neuroscience and psychiatry research is increasingly recognized and acted upon^76^, since sex differences play a role in the incidence, treatment and manifestation of psychopathology^77,78^. The precise role of sex and gender in metacognition deserves more attention and should be characterized further in future research.

Our data shows no convincing evidence for an exaggerated decrease/increase in confidence during loss/gain anticipation in OCD/GD, respectively. However, the group*incentive interaction approached significance, with increased confidence in GD patients compared to both OCD patients and HCs, specifically in the gain condition. This finding agrees with literature demonstrating increased reward sensitivity in GD^79,80^. Confidence in OCD patients has been mostly studied using metamemory paradigms, and abnormalities were most profound in OCD-relevant contexts^81–86^. Earlier studies probing confidence in GD are sparse, and whilst they all did show an effect of overconfidence in (sub)clinical problem gamblers, none of the studies actively controlled for performance differences, making it difficult to draw strong conclusions about confidence biases^16,17,87^.

Since confidence in GD and OCD did not differ from the healthy population we cannot technically speak of confidence ‘abnormalities’ in GD and OCD. Future work is necessary to study the link between compulsivity and confidence more directly. One interesting method is transdiagnostic research to study metacognition in psychiatry. Transdiagnostic research methods are useful, since (meta)cognition might relate more closely to symptoms than diagnoses, due to high levels of comorbidity and heterogeneity of symptoms within disorders. Indeed, a transdiagnostic factor of ‘anxious-depression’ was negatively related to confidence, whereas ‘compulsive behavior and intrusive thoughts’ were positively related to confidence and showed decoupling of confidence and behavior by diminished utilizing of perceptual evidence for confidence judgments^88^. This latter result is in line with our findings of diminished evidence integration into confidence judgments in GD patients.

The brain areas we found to be related to confidence and incentive processing converge with earlier work. Confidence was found to be positively related to the VMPFC via automatic processing at the choice moment^20,46,47,55^. Early certainty processing was also positively related to activity in the VS and precuneus^39,49,51^. We also observed a wide-spread network of areas negatively related to early certainty, containing the dACC, dorsolateral PFC, insula, inferior parietal lobe and midfrontal gyrus, a network repeatedly associated with uncertainty and metacognitive processes^39,44,45,51^. Also, well-known relationships between reward processing and activity in both VS and VMPFC^21,22^ were replicated. Moreover, we found negative relationships between incentive value and BOLD activity in the central executive network (i.e. lateral PFC and middle frontal gyrus), as well as superior temporal gyrus^89,90^. Confidence was found to be related to VMPFC activity, not only at choice moment, but also during rating^20,46,47^.Overall, our fMRI findings closely resemble activation patterns previously shown in healthy populations.

We also replicated the effect of incentive condition on VMPFC baseline activity and on the correlation of VMPFC activity with confidence, which was highest in gain conditions, which we also found in the VS^30^. While we found aberrant evidence integration in GD patients on a behavioral level, we did not find any group differences in evidence processing on neurobiological level. Interestingly, OCD patients showed a decreased baseline VMPFC activity during incentive/rating moment, which fits with earlier work showing neurobiological deficits in a ‘ventral motivational circuit’ including the VMPFC^91,92^. However, we did not find any interactions with incentive condition in the VMPFC activity related to either signed evidence or confidence.

In sum, contrary to our hypotheses, we did not find neurobiological deficits directly related to confidence or to the effects of incentive on confidence in our clinical samples. This might not be surprising, given that the behavioral group effects were small (and disappeared when controlling for demographics), which limited our ability a priori to find impairments in neural circuits mediating confidence processes. Because, to our knowledge, the present study represents the first attempt in investigating the joint neural basis of metacognitive and reward processes in both GD and OCD, further study - e.g. looking into transdiagnostic variations of symptoms - might be more powerful in detecting clinically useful neurocognitive signatures of those processes than the present clinical case-control comparisons^93^.

## Acknowledgements

Data collection for this work was funded by two independent personal Amsterdam Brain and Cognition (ABC) Talent grants to JL and RvH, and a NWO Veni Fellowship (grant 451-15-015) granted to ML. ML is supported by a Swiss National Fund Ambizione Grant (PZ00P3_174127) and an ERC Starting Grant (ERC-StG-948671), JL is supported by a NWO VENI Fellowship grant (916-18-119).

