## Supplementary Materials for "Metacognition and the effect of incentive motivation in two compulsive disorders: gambling disorder and obsessive-compulsive disorder"

*: Shared last authors

### Supplemental Methods

#### Participants

All subjects underwent screening with the MINI structured psychiatric interview to confirm the absence of any other psychiatric disorder[1]. OCD symptom severity was measured using the Yale-Brown Obsessive Compulsive Scale (YBOCS)[2], and GD symptom severity was measured using the Problem Gambling Severity Index (PGSI)[3]. Anxiety symptoms were assessed using the Hamilton Anxiety Rating Scale (HAMA)[4] and depression symptoms using the Hamilton Rating Scale for Depression (HDRS)[5]. We analyzed whether age, sex, IQ, Y-BOCS, PGSI, HAMA and HDRS score differed between the three groups using ANOVAs for all variables but sex, which was assessed using a Chi-square test. When appropriate, two-sample t-tests were executed post-hoc.

#### Exclusion Criteria

The exclusion criteria included having a diagnosis of major depressive disorder, bipolar disorder, psychotic disorders, substance-use disorders, using tricyclic antidepressants or antipsychotics, having any contraindications for MRI, and having a history of or current treatment for neurological disorders, major physical disorders or brain trauma.

Moreover, session-level behavioral and fMRI data were excluded when task accuracy was below 50%, which would signal below chance level performance and would hinder our interpretation of the underlying cognitive processes. Also, when subjects did not show sufficient variation in their confidence reports (standard deviation of <5 confidence points), which indicates careless responding, those data were excluded. Session-level fMRI data was additionally excluded when participants displayed more than 3.5 mm head movement in any direction. Overall, for the behavioral analyses, this led to the full exclusion of four GD patients and one OCD patient, as well as one out of two session exclusions for four GD patients, two OCD patients and two HCs. For the fMRI analyses, three additional GD patients, one OCD patient and two HCs were fully excluded, as well as one out of two session exclusions for three additional GD patients and one OCD patient.

#### Experimental Procedure

After demographic and clinical interviews, all participants performed an initial calibration session (consisting of 144 trials) to tailor the difficulty level of the task to each individual. This was done to keep average performance similar across individuals. Following, all subjects performed two fMRI sessions, each consisting of 72 trials (24 per incentive condition), presented in a random order. After the fMRI task, six random trials were drawn (i.e., two of each incentive condition) on which the final payment was based, and the total amount of points was converted to money.

**Behavioral Analyses: Properties of Confidence Judgments**

Similarly to Lebreton et al. (2018)[6], we performed additional behavioral analyses to confirm three main properties of confidence judgements, as theorized in a recent paper by Sanders and colleagues[7]. There, the authors outlined three main properties of confidence judgments, which should be observed if participants compute the probability of a choice being correct given some level of noisy evidence: (1) confidence ratings correlate with the probability of being correct; (2) the link between confidence ratings and evidence is positive for correct and negative for incorrect responses; (3) the link between evidence and performance differs between high and low confidence trials.

- To assess the first property, we sorted trials according to the confidence ratings at the individual level. Then, we averaged trials over 8 bins per participant, and computed the frequency of correct choices in each bin. Finally, the correlation between the bins' confidence and performance was computed at the individual level. These measures were positively correlated ($R = 0.59 \pm 0.03$; **Figure S1A**).
- To assess the second property, the following linear regression was estimated at the individual level, using all trials from the confidence elicitation task (Model 1):
  $Conf = \beta 0 + \beta_1 \times Correct \times Evidence + \beta_2 \times Incorrect \times Evidence$,
  where **Incorrect** is a dummy variable coding for incorrect answers, and **Correct** is a dummy variable coding for correct answers. Then, we tested the parameters of this model at the population level using one-sample t-tests. The results (**Figure S1B**), summarized in the table below (**Table S1**), demonstrate that confidence judgments are indeed positively associated with evidence for correct trials, and negatively for incorrect trials.
- To assess the third property, we proceeded similarly to the second: the following logistic regression was estimated at the individual level, using all trials (Model 2).
  $Correct = \beta 0 + \beta_1 \times High \times Evidence + \beta_2 \times Low \times Evidence$, where **High** is a dummy variable coding for high confidence trials (i.e. confidence > median(confidence)), and **Low** is a dummy variable coding for low confidence trials (i.e. confidence ≤ median(confidence)). Then, the parameters of this model were tested at the population level, using one-sample t-tests. The results (**Figure S1C**), summarized in the table below (**Table S1**), indeed demonstrate that the curve has a steeper slope in the high than in the low confidence trials, as was expected. Nota bene: when inspecting the data for the last model, we observed that the regression model was inestimable for two subjects. This was due to the median-split of the early certainty trials into high and low variants, since for both subjects the amount of low confidence trials was not

sufficient (i.e., <nbins) to estimate the model, since the distribution of their confidence judgments was very skewed. This resulted in $\boldsymbol{\beta}_2$ being inestimable. Therefore, we excluded those two subjects (only) from the analyses of the last model when testing at the population level, whilst including them for the other models.

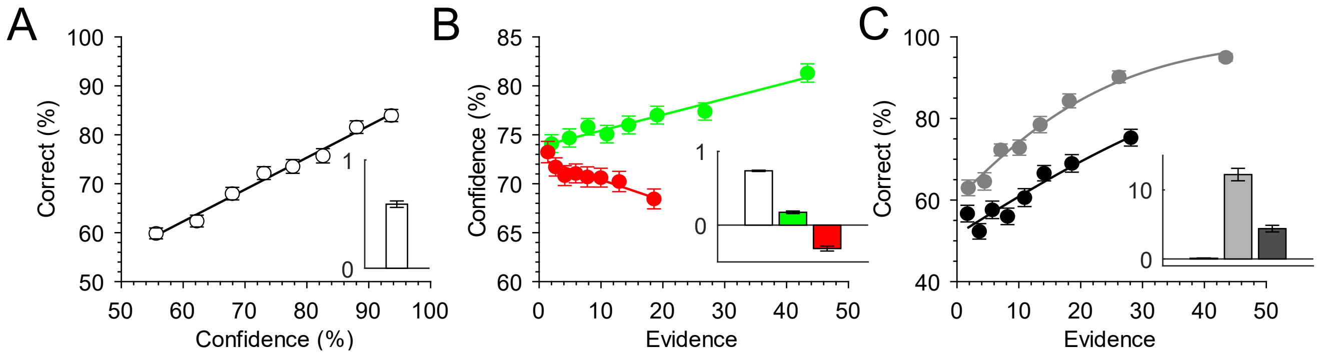


**Figure S1: Properties of Confidence Judgments**
A: observed performance (% correct choices) as a function of reported confidence. B: reported confidence as function of evidence for correct (green) and incorrect (red) choices. C: observed performance (% correct choices) as a function of evidence, for high (gray) and low (black) confidence trials. The insets presented on the side of each graph depict the results of the population-level analyses on the correlation coefficients (A) or on the regression coefficients (B and C). Error bars indicate inter-subject standard errors of the mean. *: P<.05; **: P<.01; ***P<.001

| | |
|---|---|
| **Model 1 (Figure S1B)** | |
| **Intercept ($\beta_0$)** | $\beta = 0.7366 \pm 0.0085$<br>$t_{109} = 86.9948$<br>P = 1.5197e-102 |
| **Confidence/Evidence Correct Answers ($\beta_1$)** | $\beta = 0.1736 \pm 0.0174$<br>$t_{109} = 9.9870$<br>P = 4.5659e-17 |
| **Confidence/Evidence Incorrect Answers ($\beta_2$)** | $\beta = -0.3184 \pm 0.0335$<br>$t_{109} = -9.5124$<br>P = 5.5456e-16 |
| **Model 2 (Figure S1C)** | |
| **Intercept ($\beta_0$)** | $\beta = 0.0987 \pm 0.\ 0446$<br>$t_{107} = 2.2134$<br>P = 0.0290 |
| **Performance/Evidence High confidence ($\beta_1$)** | $\beta = 12.2197 \pm 0.\ 9019$<br>$t_{107} = 13.5494$<br>P = 4.0743e-25 |
| **Performance/Evidence Low confidence ($\beta_2$)** | $\beta = 4.3919 \pm 0.\ 4810$<br>$t_{107} = 9.1305$<br>P = 4.1081e-15 |
| **Difference ($\beta_1 - \beta_2$)** | $t_{107} = 10.7705$<br>P = 7.3768e-19 |

**Table S1: Results of properties of confidence judgments**

#### Behavioral Analyses: Properties of Early Certainty

Here we provide further details about the computation and properties of the early certainty variable. To verify that our model of early certainty is an appropriate proxy of confidence judgments, we performed similar behavioral analyses to confirm the three main properties of confidence judgments still hold for our early certainty variable. We performed identical analyses, substituting subjective confidence judgments for early certainty values.

Our results show that the measures of early certainty and performance are highly correlated (R = 0.73 ± 0.0362; **Figure S2A, Table S2**). Early certainty is also positively associated with evidence for correct trials, and negatively for incorrect trials (**Figure S2B, Table S2**). Finally, the relationship between performance and evidence is indeed higher in trials with high early certainty versus low early certainty (**Figure S2C, Table S2**). Nota bene: when inspecting the data for the last model, we observed that the regression model was inestimable for four subjects. This was due to the median-split of the early certainty trials into high and low variants, where these four subjects had an average performance of 100% in the *high confidence* trials, making $\boldsymbol{\beta}_1$ inestimable. Therefore, we excluded those four subjects from the analyses of the last model when testing at the population level, whilst including them for the other models.

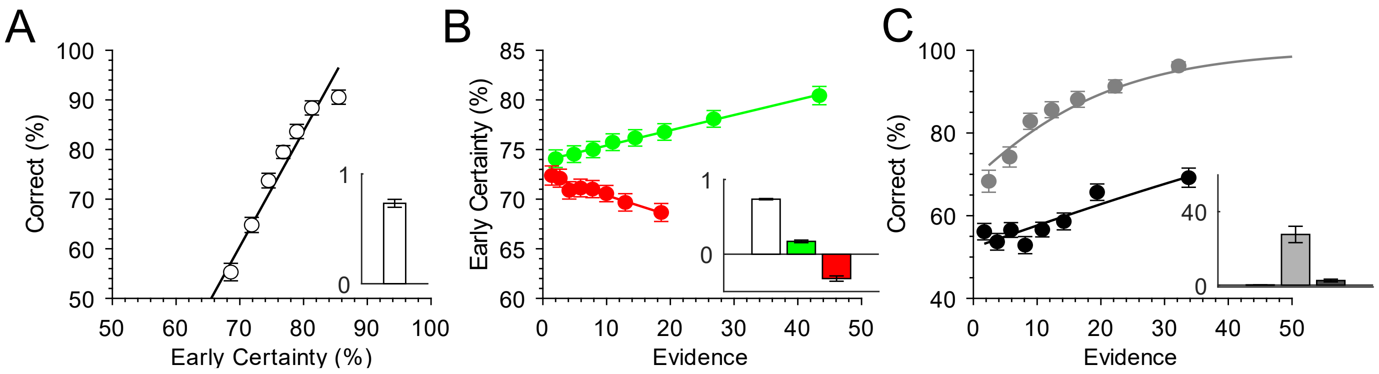


**Figure S2: Properties of Early Certainty**
A: observed performance (% correct choices) as a function of early certainty. B: early certainty as function of evidence for correct (green) and incorrect (red) choices. C: observed performance (% correct choices) as a function of evidence, for high (gray) and low (black) early certainty trials. The insets presented on the side of each graph depict the results of the population-level analyses on the correlation coefficients (A) or on the regression coefficients (B and C). Error bars indicate inter-subject standard errors of the mean. *: P<.05; **: P<.01; ***P<.001

| **Model 1 (Figure S2B)** | |
|---|---|
| **Intercept ($\beta_0$)** | $\beta$ = 0. 7364 ± .0084<br>$t_{109}$ = 87.1574<br>P = 1.2434e-102 |
| **Confidence/Evidence Correct Answers ($\beta_1$)** | $\beta$ = 0.1714 ± .00177<br>$t_{109}$ = 9.7058<br>P = 2.0064e-16 |
| **Confidence/Evidence Incorrect Answers ($\beta_2$)** | $\beta$ = -0.3266 ± .0355<br>$t_{109}$ = - 9.2108<br>P = 2.6975e-15 |
| **Model 2 (Figure S2C)** | |
| **Intercept ($\beta_0$)** | $\beta$ = 0. 0858 ± 0. 0484<br>$t_{105}$ = 1.7401<br>P = 0.0848 |
| **Performance/Evidence High confidence ($\beta_1$)** | $\beta$ = 27.5829 ± 4. 4499<br>$t_{105}$ = 6.0848<br>P = 1.9264e-08 |
| **Performance/Evidence Low confidence ($\beta_2$)** | $\beta$ = 2.5038± 0. 7493<br>$t_{105}$ = 3.2801<br>P = 0. 0014 |
| **Difference ($\beta_1$ - $\beta_2$)** | $t_{105}$ = 5.5184<br>P = 2.4830e-07 |

**Table S2: Results of properties of early certainty**

Moreover, to validate that our model of early certainty correlates highly with subjective confidence and choice and stimulus features, but does not show a statistical relationship with incentives, we built a linear mixed-effects model using the afex package in R. We used early certainty as dependent variable and added RT, accuracy, evidence and the interaction between evidence and accuracy as predictors. Indeed, the results showed that RT ($F_{1,\ 15169}$ = 11622.7231, p<.001), accuracy ($F_{1,15157}$ = 1855.0251, p<.001) and the accuracy * evidence interaction ($F_{1,15156}$ = 626.7032, p<.001) all significantly contributed to early certainty, while no effect of incentive value on early certainty was found ($F_{1,15154}$ = 1.9232, p=.1462).

**Behavioral Analyses: Model Comparisons**

We iteratively built linear mixed effects models (LMEMs) and compared those by assessing model fit by using Chi-square tests on the log-likelihood values, and by comparing of the AIC and BIC model values. We started with a basic model with fixed effects of incentive, group and their interaction on confidence, together with a random subject intercept and slope of incentive per subject. Model predictors of accuracy and evidence, together with their interaction and the interaction with group were added whenever it significantly improved model fit. See Table 1 in the main text for the model comparison results. The final model (Model 1) consisted of fixed effects of incentive, group, accuracy and evidence (z-scored) and interactions between incentive and group, as well as two-way and three-way interactions between evidence, accuracy and group. All models included trial-by-trial data, and a random subject intercept as well as a random slope of incentive per subject.

**Behavioral Analyses: Integration of Evidence in Confidence Judgments**

Theoretical models of confidence formation suggest that confidence builds – at least partly - on the integration of noisy perceptual evidence used for decision-making[7,8]. A resulting signature of confidence is its statistical dependence on an interaction of accuracy and perceptual evidence, which is typically illustrated as an 'X-pattern' where confidence increases/decreases with increasing evidence for correct/incorrect decisions, respectively. To study if GD and OCD patients show aberrant integration of evidence in confidence signals, we have included a three-way interaction term between evidence, accuracy and group in Model 1. Post-hoc testing was performed by comparing the groups on the slopes of evidence integration in confidence separately for correct and incorrect trials using the emtrends() function.

**Behavioral Analyses: Confidence Calibration**

Confidence calibration – also known as confidence bias – is the difference between average confidence and average performance per subject. If this measure is positive, this indicates overconfidence, whereas negative numbers indicate underconfidence. We calculated confidence calibration for each subject per incentive condition. We then performed a mixed ANOVA implemented in the afex package, to test for main effects of incentive conditions, groups, and their interaction. When a main effect was found significant, we performed post-hoc testing using the emmeans package, correcting for multiple comparisons using Tukey's method.

**Behavioral Analyses: Metacognitive Sensitivity**

Metacognitive sensitivity is a measure that indicates how well one's confidence judgments discriminate between one's correct and incorrect answers. One of the metrics used to express metacognitive sensitivity is *discrimination*. Discrimination is calculated as the difference between one's average confidence in their correct answers and their incorrect answers. The higher this metric, the more sensitive one's metacognitive abilities are. Another metric for sensitivity is *meta-d'*, which represents how much information in signal-to-noise units is available for the formation of confidence judgments [9]. The higher meta-d', the higher the metacognitive sensitivity.

We calculated discrimination for each subject per incentive condition. Moreover, we computed meta-d' per incentive and group using a hierarchical Bayesian framework[10]. We then performed two mixed ANOVA implemented in the afex package, to test for main effects of incentive conditions, groups, and their interaction on discrimination and meta-d' separately.

**fMRI Analyses: Acquisition & Preprocessing**

All our analyses were performed using MATLAB with SPM12 software (Wellcome Department of Cognitive Neurology, London, UK). Raw multi-echo functional scans were weighed and combined into 570 single volumes per scan session (64), using the first 30 dummy scans to calculate the optimal weighting of echo times for each voxel by applying a PAID-weight algorithm. During the combining process, realignment was performed on the functional data by using linear interpolation to the first volume. Subsequently, the functional images were co-registered, segmented for normalization to MNI space and smoothed. To reduce motion-related artifacts, the Art-Repair toolbox[11] was used to detect large volume-to-volume movement and repair outlier volumes. Outliers were detected using a threshold for the variation of the mean intensity of the BOLD signal and a volume-to-volume motion threshold. A threshold of 1.5% variation from the mean intensity was used to detect and repair volume outliers by interpolating from the adjacent volumes.

**fMRI Analyses: General Linear Models**

GLM 1 consisted of three regressors for each timepoint: 'choice', 'incentive/rating' and 'feedback', to which parametric modulators (pmods) were added. All regressors were specified as stick functions time-locked to the onset of the respective events. The choice regressor was modulated by two pmods: early certainty (z-scored on subject level) and button press (left or right) to control for motor-related activation. The incentive/rating regressor was modulated by two pmods: incentive value ([-1,0,1]) and confidence rating (z-scored on subject level). The feedback regressor was additionally modulated by a pmod representing choice accuracy.

GLM 2 consisted of regressors for each of two time points (choice moment and incentive/rating moment) and three incentive conditions, as well as a single regressor at feedback moment, resulting in a total of seven regressors. All regressors at choice moment were modulated by a pmod of button press (left/right) and signed evidence: a variable that signifies the interaction between evidence and accuracy. Signed evidence was calculated as the absolute value of evidence in case of correct answers and the negative absolute evidence (i.e. –abs(evidence)) in case of incorrect answers. All regressors at rating moment were modulated by a pmod of confidence, and the feedback regressor was modulated by a pmod of accuracy. Thus, for all these events we could examine both baseline activity and regression slopes relating to their respective pmod.

For both GLMs pmods were not orthogonalized and thus competed to explain variance. We included six motion parameters as nuisance regressors. Regressors were modeled separately for each scanning session and constants were included to account for between-session differences in mean activation. All events were modeled by convolving a series of delta functions with the canonical hemodynamic response function (HRF) at the onset of each event and were linearly regressed onto the functional BOLD-response signal. Low frequency noise was filtered with a high pass filter with a cut off of 128 seconds. We controlled for the number of sessions while making the first-level contrasts. All contrasts were computed at subject level and then taken to group level analyses. For GLM 1 we assessed group differences by performing a one-way ANOVA to our contrasts of interest, using an F-contrast test to test for any group differences (i.e. [1 -1 0; 0 1 -1]). In addition, to gain a complete picture of areas involved in our contrasts of interest, we grouped all subjects together and performed one-sample t-tests against 0.

### Supplemental Results

#### Demographics

Age was not significantly different between the three groups (F2,107 = 0.253, p > 0.75), but IQ was, (F2,107 = 3.222, p = 0.0438). Post-hoc t-tests showed that HC subjects had a significantly higher IQ score than GD patients (t = 2.53, p=0.014). As expected, Y-BOCS scores and PGSI scores differed significantly between groups (F2,107 = 322.2, p<.001 ;F2,107 = 380.5, p<.001, respectively), with OCD patients having higher Y-BOCS scores than HCs (t = -16.97, p<.001) and GD patients (t = -36.67, p<.001), and GD patients having higher PGSI scores than HCs (t = -15.99, p<.001) and OCD patients (t = -14.32, p<.001). HAMA scores were significantly different between groups (F2,107 = 48.02, p<.001), post-hoc tests revealed higher HAMA scores for OCD patients than HCs (t = -8.50, p<.001) and GD patients (t = 4.58, p<.001), and higher HAMA scores for GD patients compared to HCs (t = -2.44, p=.002). HDRS scores were significantly different between groups (F2,107 = 24.97, p<.001), with higher scores for OCD versus HC (t = -7.76, p<.001), and higher scores for GD versus HC (t=-3.03, p=.005). Lastly, using a Chi-square test we found a significant difference in sex distribution between the groups (X = 14.483, df = 2, p<.001),

#### Behavioral Descriptive Results

Here we show the descriptive results that are depicted in **Figure 2**.

| Group | Incentive | Confidence | Accuracy | RT | Evidence |
|---|---|---|---|---|---|
| GD | Loss | 76.31 +- 1.91 | 71.22 +- 1.66 | 1175.57 +- 81.19 | 15.05 +- 1.00 |
| GD | Neutral | 78.56 +-1.64 | 73.07 +- 1.61 | 1135.56 +- 71.15 | 17.60 +- 1.14 |
| GD | Gain | 81.12 +-1.84 | 71.37 +- 1.69 | 1132.38 +- 79.91 | 16.97 +- 1.26 |
| OCD | Loss | 71.30 +- 1.85 | 72.62 +- 1.42 | 1215.41 +- 74.70 | 13.84 +- 0.79 |
| OCD | Neutral | 73.27 +- 1.70 | 73.14 +- 1.39 | 1219.32 +- 65.71 | 15.56 +- 0.94 |
| OCD | Gain | 73.70 +- 1.65 | 75.07 +- 1.78 | 1224.67 +- 68.70 | 14.69 +- 0.84 |
| HC | Loss | 73.02 +- 1.03 | 70.87 +- 1.27 | 1130.16 +- 44.63 | 14.87 +- 0.63 |

| | | | | | |
|---|---|---|---|---|---|
| HC | Neutral | 75.05 +- 1.01 | 71.86 +- 1.19 | 1142.39 +- 48.26 | 16.90 +- 0.79 |
| HC | Gain | 75.68 +- 1.07 | 72.35 +- 1.27 | 1149.16 +- 46.71 | 16.05 +- 0.76 |

**Table S3: Descriptive behavioral results**. Shown here are the descriptive results of confidence, accuracy, reaction times (RT) and evidence per group and incentive condition. Shown are means +- sems.

**Behavioral Analyses: Integration of Evidence in Confidence Judgments**

The evidence integration effect differed per group, as signaled by a significant three-way interaction between accuracy, evidence and group (F2,15094 = 3.0533, p=0.04723) (Figure 3, Supplementary Table 3). Post-hoc, we compared the groups on the slopes of evidence integration in confidence separately for correct and incorrect trials using the emtrends() function, and found that the slope for evidence integration into confidence was less steep for correct answers in GD patients compared to both HCs (GD - HC = -1.712 +- 0.283, Z-ratio = -6.057, p<0.001) and OCD patients (GD - OCD = -2.110 +- 0.357, Z-ratio = -5.912, p<0.001). This indicates that GD patients' confidence ratings were less influenced by the perceptual evidence when they made a correct choice. No differences between OCD patients and HC were found regarding evidence integration effects.

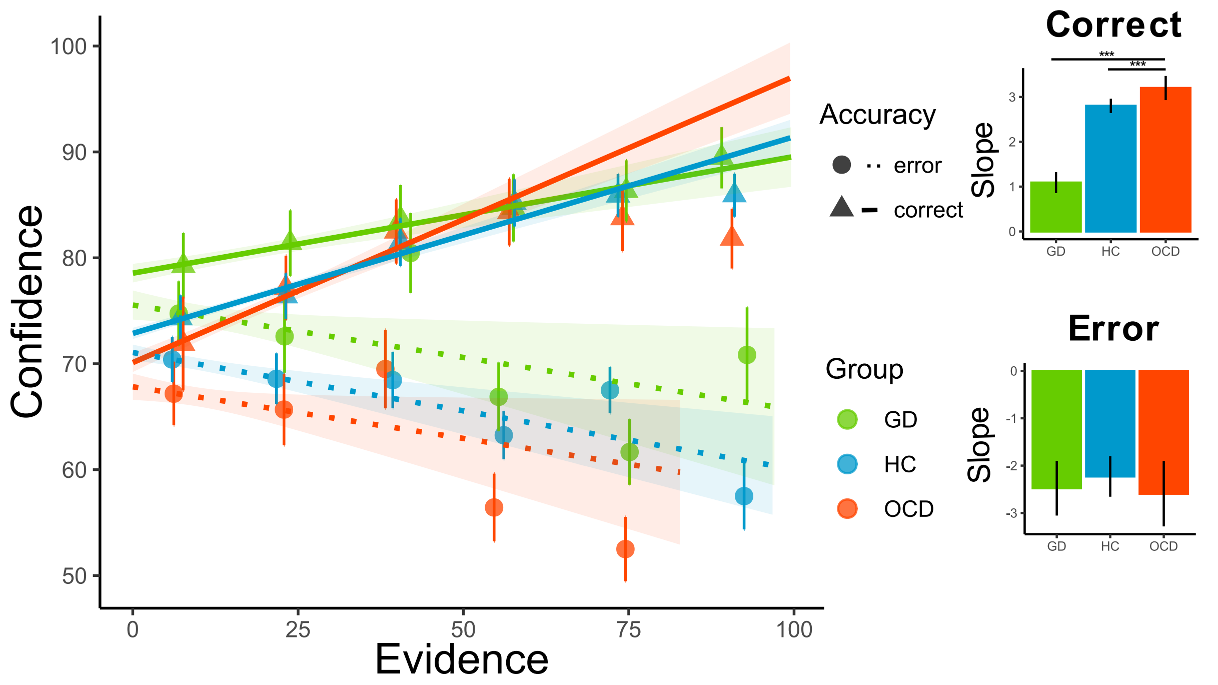


**Figure S3: Evidence integration per group and accuracy level in confidence.** Linking evidence, accuracy and group. Triangles represent mean reported confidence as a function of evidence for correct answers, and dots for incorrect answers, with different colors for the three groups. The solid lines represent the best linear regression fit for each group separately at the population level for correct answers, and the dotted lines for incorrect answers. Error bars represent SEM per group, shaded areas represent 95% confidence interval. Insets represent slopes (estimated marginal means of trends, taken from emtrends() function, error bars represent SEM) of correct and incorrect answers per group. Results from post-hoc testing are shown, where the slope for correct answer is significantly lower for gambling disorder (GD) versus both healthy controls (HC) and obsessive-compulsive disorder (OCD) (* p<0.05, ** p<0.01, *** p<0.001).

**Behavioral Analyses: Confidence Calibration**

We found a significant main effect of group (F(2,107)=4.40, p = 0.015), but no effect of incentive, nor an interaction effect between group and incentive. Post-hoc tests showed that GD patients showed increased calibration compared to OCD patients ($t_{107}$: -2.967, p = 0.0103), but no differences between GD or OCD patients and HC subjects. This indicated that GD patients are more overconfident

**Behavioral Analyses: Metacognitive Sensitivity**

We did not find a significant main effect of group or incentive, nor an interaction effect between group and incentive, both for discrimination and meta-d'. Average discrimination values were positive and average meta-d' was close to 1, indicating sensitive metacognition.

**Behavioral Analyses: Clinical Correlations**

We performed additional correlational analyses to explore whether subject's mean confidence level correlates with various clinical questionnaires of interest, separately for OCD and GD patients. In OCD patients there were no significant correlations with severity of OCD symptoms as measured with the YBOCS (p>.5) or with obsessive beliefs measured with the OBQ-44 (p>.5). In GD patients there was also no significant correlation with symptom severity measured using PGSI (p>.4), but there was a significant positive correlation between confidence level and BAS (Behavioral Approach System) scores (r = 0.4608, p = 0.01784).

**Behavioral Analyses: Evidence Across Conditions**

Due to a technical bug, perceptual evidence was not equal across incentive conditions. We performed a mixed ANOVA with within-subject factor incentive and between-subject factor group, which showed that evidence differed significantly over incentive conditions (F2,205=39.94 p<.001), but not over groups (F2,107=0.94 p>.3), and no interaction between incentive and group was found (F3.83,205=0.82 p>.5). Post-hoc testing using t-tests revealed that evidence was highest in neutral, followed by gain, followed by loss condition (neutral versus loss: t-ratio= 7.844, p<.001; neutral versus gain: t-ratio=3.306, p=0.001; gain versus loss: t-ratio: 5.537, p<.001). Since evidence did not differ between the groups, it cannot account for any group differences we find in our data. Importantly, there are no effects of incentive on performance. Moreover, the difference in evidence over incentive conditions does not drive our incentive-induced confidence bias, since we do find a parametrically increase in confidence over incentive value, with a significant difference between all pairs. This means that confidence is higher in gain versus neutral conditions, even though evidence was significantly higher in neutral versus gain conditions. This shows that even though trials were easier in the neutral condition, participants were still more confident when they could gain points.

##### Behavioral Analyses: Clinical Groups With Their Own Control Group

In order to explore whether the behavioral analyses as in the main results with better matched control groups to the demographics of the two clinical groups would reveal similar results, we selected two subsets from our bigger sample of HCs (OCD control group N = 31, GD control group N = 32, with a slight overlap of N=8) of control groups to compare them with the two clinical groups.

Even though the groups were better matched and did not significantly differ from the clinical groups on terms of age, sex or IQ, we found similar results. When comparing OCD patients to a matched HC group, using Model 4 as described in the methods, we only found a significant main effect of incentive ($F_{2,111} = 9.5665$, $p<.001$), accuracy ($F_{1,8217} = 337.5033$, $p<.001$), evidence ($F_{1,8214} = 6.7033$, $p=.009$), an interaction effect between accuracy and evidence ($F_{1,8224} = 118.2244$, $p<.001$) and an interaction effect between group and accuracy ($F_{1,8227} = 7.9859$, $p=.004726$). No group effects were found.

When comparing GD patients to a matched HC group, using Model 4 from the main methods, we found significant main effects of incentive ($F_{2,60} = 15.6065$, $p<.001$), accuracy ($F_{1,8033} = 365.6563$, $p<.001$), evidence ($F_{1,8029} = 7.7733$, $p=.0053$), an interaction between accuracy and evidence ($F_{1,8027} = 117.9345$, $p<.001$), and between accuracy evidence and group ($F_{1,8027} = 6.5439$, $p=.0105$). No main group effect was found. Post-hoc analyses showed that that the slope for evidence integration into confidence is less steep for correct answers in GD patients compared to HCs (GD - HC = -1.329 +- 0.326, Z-ratio = -4.071, $p<0.001$).

These additional analyses thus show that even when using a better matched control group, we find no evidence for abnormalities in confidence level for OCD nor GD patients. For GD patients, we do, however, replicate that GD patients have a lower slope of evidence integration in confidence for correct answers compared to HCs.

##### fMRI: Interaction Between Metacognition and Incentives in VS (GLM 2)

We performed an ROI analysis by leveraging our factorial design. We extracted VS activations for both time points (choice and rating), all incentives (loss, neutral and gain), all groups (HC, OCD and GD), for both baseline activity and a regression slope with (1) signed evidence and (2) confidence judgments for all these events.

First, one-sample t-tests showed that, overall, VS baseline activations did not differ from 0 at choice moment (t100 = -0.317, p >0.75), while it was positive for baseline activations at rating moment (t100 = 8.238, p < 0.001). The correlations between VS activity and signed evidence at choice moment was significantly positive (t100 = 4.985, p < 0.001). However, the correlation between VS activity and confidence at rating moment did not differ from 0 (t100 = 1.664, p = 0.099) (**Figure S3**). This implies that activity in VS is related to incentive presentation, but also that it is related to signed evidence (i.e. the interaction between accuracy and evidence, showing that VS activity was lowest when one had high levels of evidence but was incorrect, and highest when one had a lot of evidence and was in fact correct). Then, we turned to see whether there were effects of incentive condition and group around this general signal. As expected, at choice moment there were no effects of incentive condition on VS baseline activity, nor on its correlation with the signed evidence signal (i.e. slope) (**Figure S3, Table S4**). Moreover, we did not find a group nor an interaction effect on both baseline VS activity and the correlation with signed evidence at choice moment. At rating moment, however, incentive condition had a significant effect on both the baseline VS activity, as well as its correlation with confidence. Post-hoc testing showed that the baseline VS activity was highest during gain, followed by loss, and lowest during neutral (loss versus gain: t196: -4.590, p < 0.001, neutral versus gain: t196: -7.710, p < 0.001, loss versus neutral: t196 = 3.119, p = 0.006). The correlation of VS activity with confidence was significantly higher (i.e. increased slope) in gain versus neutral (t196 = -2.607, p = 0.0265), while no differences between gain and loss, or between neutral and loss were found. Moreover, there was a significant group effect on VS baseline activity during rating moment. This effect did not remain significant in the post-hoc tests, however, which showed that GD subjects had subthreshold decreased activity compared with HCs, averaged over incentive conditions (t98 = -2.272, p = 0.0646). No interaction effects between group and incentive were found on baseline activity or its correlation with confidence at rating moment.

| | **Incentive** | **Group** | **Incentive:Group** |
|---|---|---|---|
| **Choice Baseline** | F(1.92, 188.46) = 0.16, p = 0.846 | F(2, 98) = 1.32, p = 0.271 | F(3.85, 188.46) = 0.66, p = 0.615 |
| **Choice Slope ‘Signed Evidence’** | F(1.99, 195.35) = 1.63, p = 0.198 | F(2, 98) = 0.44, p=0.647 | F(3.98, 195.35) = 0.49, p = 0.741 |
| **Rating Baseline** | F(1.85, 181.48) = 30.08, p<**0.001** | F(2, 98) = 3.48, p = **0.035** | F(3.70, 181.48) = 0.90, p = 0.460 |
| **Rating Slope ‘Confidence Judgment’** | F(1.92, 188.55) = 3.41, p = **0.037** | F(2, 98) = 1.68, p = 0.192 | F(3.83, 188.55) = 0.69, p = 0.593 |

**Table S4: Results of VS ROI analysis.** Shown here are the results of the mixed ANOVAs of t-statistics in the ventral striatum (VS) region of interest (ROI). Shown are the main effects of incentive condition, group and their interaction effect on the choice and rating time points, focusing on both the baseline activity as well as the slope of signed evidence and confidence judgments, respectively. F-values, with corresponding degrees of freedom and p-values are reported.

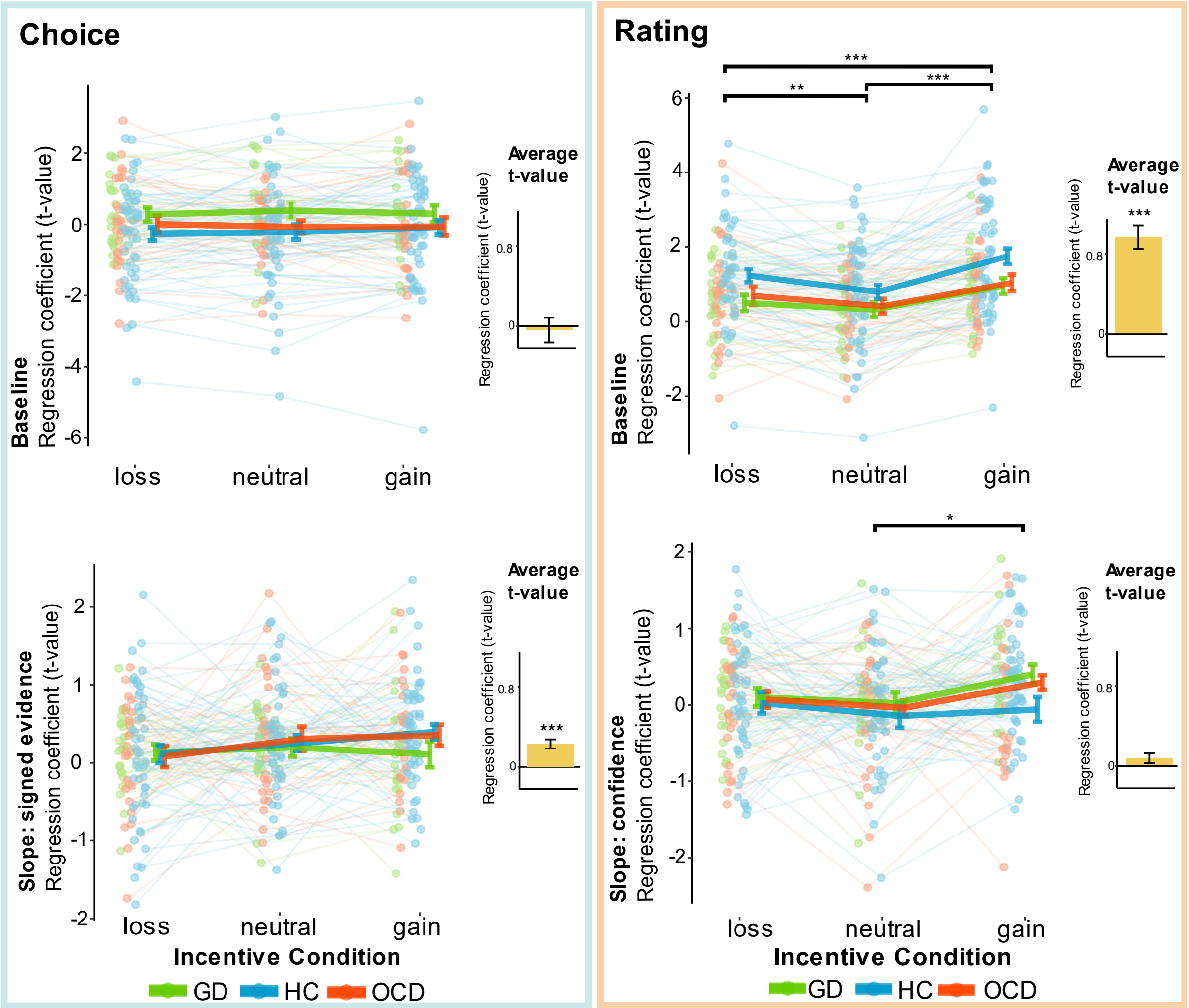


**Figure S4: Ventral striatum cortex region of interest (ROI) analysis.** T-values corresponding to baseline and regression slopes were extracted for all three groups and three incentive conditions, at two time points of interest: choice and incentive/rating moment. Green dots and lines represent gambling disorder patients, blue dots and lines represent healthy controls and red dots and lines represent obsessive-compulsive disorder patients. Dots represent individual t-statistics, and error bars represent sample mean ± SEM per group. Black bars represent significant post-hoc tests. Yellow bars represent average t-values, with corresponding significance level of one-sample t-tests against 0. (* p<0.05, ** p<0.01, *** p<0.001). GD = gambling disorder, HC = healthy control, OCD = obsessive-compulsive disorder.

11 Mazaika, Whitfield-Gabrieli, Reiss… A. Artifact repair for fMRI data from high motion clinical subjects. *Annu Meet …* 2007; : 321.
